# FlyVISTA, an Integrated Machine Learning Platform for Deep Phenotyping of Sleep in *Drosophila*

**DOI:** 10.1101/2023.10.30.564733

**Authors:** Mehmet F. Keleş, Ali Osman Berk Sapci, Casey Brody, Isabelle Palmer, Christin Le, Öznur Taştan, Sündüz Keleş, Mark N. Wu

**Affiliations:** Department of Neurology, Johns Hopkins University, Baltimore, MD 21205, USA; Department of Computer Science, Sabanci University, Tuzla, Istanbul, 34956, Turkey; Department of Biostatistics and Medical Informatics, University of Wisconsin-Madison, Madison, WI 53706, USA; Department of Neuroscience, Johns Hopkins University, Baltimore, MD 21287, USA

## Abstract

Animal behavior depends on internal state. While subtle movements can signify significant changes in internal state, computational methods for analyzing these “microbehaviors” are lacking. Here, we present FlyVISTA, a machine-learning platform to characterize microbehaviors in freely-moving flies, which we use to perform deep phenotyping of sleep. This platform comprises a high-resolution closed-loop video imaging system, coupled with a deep-learning network to annotate 35 body parts, and a computational pipeline to extract behaviors from high-dimensional data. FlyVISTA reveals the distinct spatiotemporal dynamics of sleep-associated microbehaviors in flies. We further show that stimulation of dorsal fan-shaped body neurons induces micromovements, not sleep, whereas activating R5 ring neurons triggers rhythmic proboscis extension followed by persistent sleep. Importantly, we identify a novel microbehavior (“haltere switch”) exclusively seen during quiescence that indicates a deeper sleep stage. These findings enable the rigorous analysis of sleep in *Drosophila* and set the stage for computational analyses of microbehaviors.

## Introduction

A fundamental function of the brain is to integrate information about internal states and the external environment to produce appropriate behaviors (*1*). To gain insights into this process, scientists have sought to develop methods for quantifying different aspects of animal behavior. Recent advances in computing power and machine learning have led to the rise of “computational ethology,” a field that seeks to perform automated quantification of the spatiotemporal structure of behavior (*2–5*). While animal behaviors exist across a broad swath of the 4D behavioral state-space, the vast majority of studies in this field have examined behaviors that involve large movements over short timescales.

However, for >150 years, scientists such as Darwin and Tinbergen have recognized that even subtle body part movements or small changes in posture in cats and gulls can reveal important changes in internal states such as arousal, fear, or anxiety (*6*, *7*). Despite their importance, the computational analysis of such “microbehaviors” is challenging, owing to the difficulty of reliably detecting small movements and the computationally-intense analysis of high-resolution images over long timescales.

To address this challenge, we developed FlyVISTA (Fly <u>V</u>ideo Imaging <u>S</u>ystem with <u>T</u>racking and <u>A</u>nalysis), a machine learning platform for characterizing and quantifying microbehaviors in freely moving *Drosophila*. FlyVISTA comprises a high-resolution video imaging system coupled with closed-loop perturbation using an IR laser. This system utilizes a trained deep learning pose estimation algorithm (*8*) to label 35 body parts, followed by a newly-developed computational pipeline to extract meaningful behaviors from these annotations across time.

Here, we use FlyVISTA to deeply phenotype sleep in flies. Sleep is an essential, conserved behavior (*9–17*), whose function remains unknown (*18–23*). While sleep has been most extensively studied in mammals, our understanding of the function(s) of sleep would be greatly aided by rigorous analyses of sleep in simple genetically-tractable non-mammalian organisms (*9*, *11*, *24–26*). In addition to deepening our understanding of the molecular and cellular processes underlying sleep, such studies could reveal aspects of sleep that are conserved across evolutionarily distant species and thus likely important for its function.

Current methods for phenotyping sleep in *Drosophila* typically use single beam breaks in locomotor tubes and are relatively crude (*27*, *28*), although refinements to this approach have been made using video tracking in these tubes (*26*, *29*, *30*). We sought to develop a system to more rigorously examine sleep in freely moving animals in this species. While sleep is a broadly quiescent state, it can be characterized at the behavioral level by different microbehaviors. For example, in mammals, greater immobility or rapid horizontal eye movements are associated with deeper, more refreshing sleep and rapid eye-movement (REM) sleep, respectively (*31*, *32*). Moreover, classical observational studies of bees and cockroaches have demonstrated that sleep in these insects was associated with small changes in posture or antenna position (*14*, *33*, *34*).

We utilized FlyVISTA to characterize microbehaviors occurring during wakefulness and sleep in vinegar flies. Our findings reveal a rich repertoire of microbehaviors occurring during fly sleep, including postural relaxation, antennal drooping, oscillatory movements, and rhythmic proboscis extension (PE) (*35*), and delineate the distinct spatiotemporal dynamics of these microbehaviors. To demonstrate the utility of our system, we analyze the effects of activating two neural circuits commonly used to promote sleep in *Drosophila*. These experiments reveal that optogenetic activation using a driver labeling dorsal fan shaped body (dFB) neurons (*36*) promotes micromovements, not sleep, whereas activating R5 neurons (*37*, *38*) induces PEs followed by persistent sleep. Notably, we identify a novel microbehavior termed “haltere switch” (HS), which involves a discrete movement of the halteres, organs that are functionally analogous to the vestibular system in mammals (*39*, *40*). Unlike PEs, this HS behavior is only seen during quiescence, but similarly defines a deeper stage of sleep in *Drosophila*. Together, this work reveals a novel sleep-related microbehavior and establishes a platform for the rigorous analysis of sleep and other microbehaviors in *Drosophila*.

## Results

### High-resolution video imaging and annotation of quiescent microbehaviors in freely moving *Drosophila*

Sleep can be defined by behavioral criteria as consolidated, reversible quiescence that a) preferentially occurs during a specific time of day, b) is associated with an elevated arousal threshold, and c) is under homeostatic control (*24*, *25*, *41*, *42*). To quantify sleep in *Drosophila*, most studies have relied on a definition of sleep as behavioral inactivity lasting ≥5 min (*24*) (defined by absence of a single beam-break in a thin glass tube), which is associated with an increased arousal threshold. Although convenient, there are several disadvantages to this approach. First, short bouts of locomotion could be missed using such an approach. Second, this assay misses microbehaviors in largely quiescent flies (e.g., grooming or feeding). Third, confining flies within a thin, narrow tube could induce stress or unnatural behaviors. To more rigorously characterize sleep in *Drosophila*, we sought to develop a system that combines high-resolution imaging with automated annotation and extraction of behaviors, which we have named FlyVISTA.

To facilitate the observation of natural sleep behavior, we assessed sleep in freely moving flies. In addition, a side-view imaging approach would be beneficial, because prior work in cockroaches and bees revealed that sleep in these insects was associated with specific changes in posture and body part position (e.g., drooping antennae) best viewed from the side (*14*, *33*). Thus, our high-resolution imaging setup (Fig. 1A) uses a chamber where the fly is able to turn and walk freely and is imaged from the side-view (see Methods). In this setup, the fly image is at least ∼40x larger (corresponding to ∼8 µm/pixel), compared to images from previously described fly sleep video analyses (*43*); this increased resolution enables the observation and annotation of relatively small body parts.

**Fig. 1.**
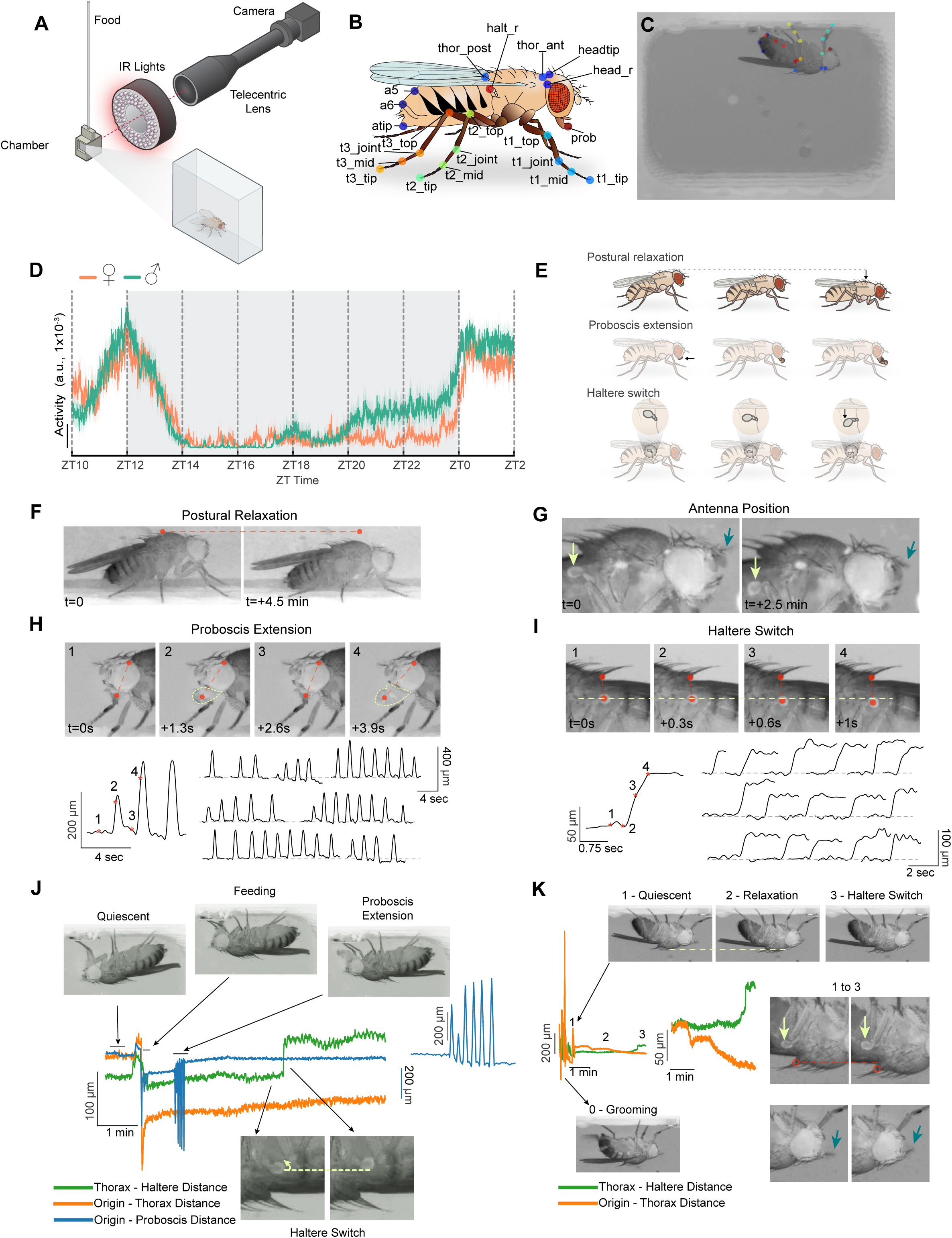
A high-resolution method to investigate quiescent behaviors in flies. (**A**) Schematic of the behavioral setup in which a fly is placed in a 7.1 (W) x 4.9 (H) x 2.8 mm (D) 3D-printed chamber with access to a liquid food capillary and imaged at high-resolution (∼8 μm / pixel). (**B**) Schematic of a fly displaying the target points tracked using DeepLabCut. 21 distinct points (35 in total when including symmetric body parts) are shown. (**C**) An example image showing a fly in the behavioral chamber with tracked body parts. (**D**) Average activity data (a.u.) per 1 min bins for male (n=23, green) and female (n=19, orange) flies from ZT10 to ZT2. Activity is based on the sum of the change in each frame for computed features derived from tracked body parts (see Methods). (**E**) Schematic illustrating microbehaviors seen during sleep. (**F**) Representative images showing the progressive lowering of the thorax position (postural relaxation) during prolonged quiescence. Dashed line connects the same pixel point across the two images. (**G**) Representative images showing downward movement of the antenna during prolonged quiescence (blue arrow). Light green arrow shows an accompanying change in haltere position. (**H**) Proboscis extension (PE) behavior is shown in 4 consecutive images (above). Red dashed line connects two points tracked by DeepLabCut: the tip of the proboscis and the dorsal edge of the eye. Bottom left: plot showing distance between the two tracked points across time; numbers/asterisks labeled on the plot correspond to the images shown. Bottom right: multiple examples of PE bouts are shown across the night. Plots show the distance between the two tracked points. (**I**) Representative images showing movement of haltere in the ventral direction during prolonged quiescence. Vertical dashed line connects the tracked points on the thorax and haltere, and horizontal dashed line indicate the same pixel points across the four images. Bottom left: plot showing distance between the 2 tracked points across time; numbers/asterisks labeled on the plot correspond to the images shown. Bottom right: multiple examples of HS behavior, shown as plots of the distance between thorax and haltere points across time. Time relative to the first image (t) is shown for (**F**-**I**). (**J** and **K**) Representative quiescence bouts exhibiting distinct behaviors with varying spatiotemporal structure. Static images of quiescence, feeding, PE, and HS (**J**) and grooming, quiescence, postural relaxation, and HS (**K**) are shown, with horizontal dashed lines connecting the same pixel points across images. Green, orange, and blue lines indicate the distance between thorax and haltere, origin (fixed point at 0, 0) and thorax, and origin and proboscis tip, respectively. Expanded traces for PE (**J**) and HS (**K**) are also shown to highlight spatiotemporal structure. In (**K**), yellow arrows point to halteres, magenta arrows point to antenna, and red circles indicate thorax.

We next trained a model using DeepLabCut (*8*) to reliably label 35 points on the fly body (Fig. 1, B and C, see Methods). Then, we generated “features” from these individual body parts (e.g., distance of proboscis tip to head), which allowed us to quantify and visualize how different body parts moved relative to each other over time. We performed video recordings of individual flies in this high-resolution setup from ZT10-ZT2 (Zeitgeber Time 10 - Zeitgeber Time 2) and first examined total locomotor activity as assessed by movement of all body parts in 1 min bins. As expected, prominent peaks of locomotor activity are observed at ZT12 and ZT0, corresponding to evening and morning peaks of activity (*44*). Prominent behavioral inactivity was observed in the early night, with gradually increasing activity in the late night, which was particularly pronounced in males (Fig. 1D) (*45*).

Several distinct microbehaviors (postural relaxation, antennal droop, proboscis extension, and haltere switch) were observed during prolonged inactivity (Fig. 1, E to I). After becoming stationary, a fly’s body posture (measured by thorax position) would sometimes gradually relax in a gravity-dependent manner (Fig. 1, F and K). It should be noted, that postural relaxation did not always precede prolonged periods of behavioral quiescence. Following this postural relaxation, the antennae of flies could be seen to droop, particularly the arista; however, the antennae of vinegar flies are small, making it difficult to consistently visualize them (Fig. 1G).

A recent study using a tethered preparation described rhythmic proboscis extension (PE) occurring during sleep in *Drosophil*a, which was associated with increased clearance of hemolymph (*35*). We also observed these rhythmic PEs during inactive periods in our freely moving flies (video S1). These PE episodes exhibit a stereotypical spatiotemporal structure, with variable duration (Fig. 1H). While PE bouts follow postural relaxation, they can occur either before or after HS/antennal droop behaviors (Fig. 1J and fig. S1A). Of note, PEs are not exclusive to periods of behavioral quiescence and are also seen during grooming (*46*), walking (*46*) or flying (*47*).

After postural relaxation, and often in synchrony with the antennae drooping, a “switch-like” movement of the halteres (vestibular-like mechanosensory organs that are sensitive to inertial forces experienced during flight) (*39*, *40*) can be seen (Fig. 1, I to K). This quiescence-associated haltere switch (HS) behavior involved a downward (i.e., towards the ventral aspect of the fly) movement that persists for ∼2-20 mins (video S2). This HS microbehavior has not, to our knowledge, been previously described in *Drosophila* or any other Diptera.

Careful observation of quiescent flies also revealed fine, fast rhythmic movements in the legs that were often present if a leg was suspended in the air (fig. S1, B and C).

These movements range between ∼1-3 Hz in frequency both within and between individual animals and have also not been previously characterized. Based on our unpublished findings in a different study, these movements likely reflect the contractions of the accessory hearts found in the leg (“leg hearts”) similar to those found in other insects (*48*). Together, high-resolution imaging and annotation reveal both known and novel microbehaviors in quiescent flies.

### Closed-loop arousal threshold analyses of *Drosophila* sleep behavior

A key aspect of the behavioral definition of sleep is the presence of an increased arousal threshold (*41*, *42*). Thus, to demonstrate that the periods of prolonged inactivity observed in our imaging setup represent sleep, we next probed arousal threshold levels. To precisely quantify arousal threshold, we used an 1064 nm IR laser whose wavelength is not detected by fly photoreceptors (*49*) to incrementally heat the fly in a closed-loop system. Fly video images were analyzed by custom software that estimated motion by using background subtraction to identify behaviors as “inactive” (no active locomotion) or “moving/arousal” (see Methods). At ≥30 min intervals, when 30 sec of “inactive” behavior was detected, the laser was activated, and its power ramped up gradually until the fly exhibited >3 sec of “moving/arousal” behavior (Fig. 2A and fig. S2A). Trials were performed from ZT10 to ZT24 (Fig. 2B). Because the absolute level of laser power required to arouse inactive flies varied substantially between individual animals, we standardized arousal threshold for each individual fly and generated a Z-score, in order to compare arousal threshold levels between animals.

**Fig. 2.**
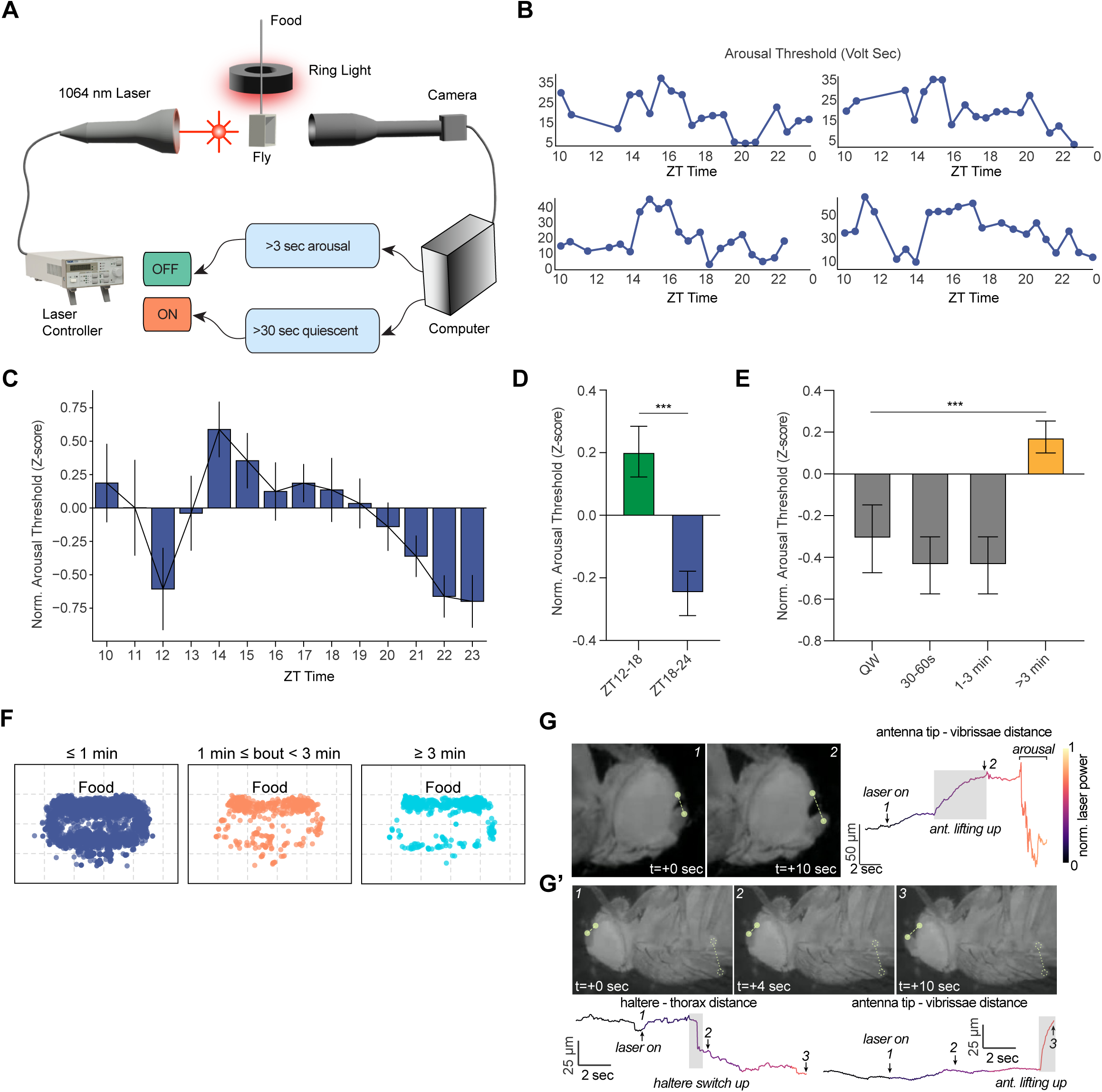
Closed-loop analyses reveal dynamic changes of arousal threshold across time. (**A**) Schematic of the closed-loop setup to test arousal threshold where a fly in chamber is placed between an IR laser and a camera. The laser is turned on after 30 sec of quiescence, and laser voltage is gradually increased over the subsequent 30 sec (fig. S2A). Once the fly exhibits 3 sec of persistent movement, the laser is turned off. (**B**) Representative plots of arousal thresholds (volt-sec) for individual female flies from four different experiments across ZT time. (**C**) Normalized arousal thresholds (Z-score) for individual perturbation bouts plotted in 60 min bins across ZT10-ZT23, showing dynamic changes of arousal threshold across time for female flies (n=16). Arousal thresholds for each fly were normalized to the mean arousal threshold for that individual fly. Error represents SEM. (**D**) Normalized arousal threshold plotted for female flies during ZT12-18 or ZT18-24 windows. Data were obtained from the same flies in (**C**). Error denotes SEM; unpaired t-test. ***, *P*<0.001. (**E**) Normalized arousal thresholds for female flies for quiet wakefulness/QW (30 sec period of locomotor inactivity prior to “laser on” accompanied by an awake “microbehavior,” i.e. feeding, grooming, defecation) (n=41 bouts), 30-60 sec quiescence prior to “laser on” (n=27 bouts), 1-3 min quiescence prior to “laser on” (n=27 bouts), or >3 min quiescence prior to “laser on” (n=137 bouts). Error denotes SEM; one-way ANOVA with post-hoc Tukey. ***, *P*<0.001. (**F**) Plots showing location of the thorax for quiescence bouts separated according to whether bouts were <1 min (blue), between 1-3 min (orange), or ≥3 min (teal). Flies spent >1 min quiescent bouts near food (n=22 flies, data are pooled from males and females). (**G**) Representative static images of changes in sleep-associated microbehaviors as laser power is increased. Top panels show antennal movement during the laser ramp-up. Representative images (top left) from time points marked on the time series data (top right). Time series data show the change in the antenna-vibrissae distance (points are shown in the top left panels) as the laser power is increased. (**G’**) Upper panels show representative static images of a HS “up” event followed by an antennal upward movement. Time series data of haltere-thorax and antenna tip-vibrissae distances are shown in lower panels, where the numbers correspond to the static images. Shaded boxes denote HS “up” and antenna “up” events. Color indicates the change in the laser power.

We first examined whether arousal thresholds correlated with how long the fly was inactive prior to the laser stimulation (fig. S2B). We found a significant, but weak, positive correlation between normalized arousal threshold and inactive bout duration.

We next asked whether arousal threshold varied across the night. We plotted arousal thresholds in 1 hr bins and found that arousal threshold was highly dynamic from ZT10-24. Arousal threshold was low at ZT12 and then peaked at ZT14, gradually falling throughout the night until reaching a nadir at ZT23 (Fig. 2C). Overall, normalized arousal threshold was significantly elevated in the first half of the night (ZT12-18) vs the second half of the night (ZT18-24) (Fig. 2D). These dynamic changes in arousal threshold could simply be due to differences in inactive bout duration throughout the night. To examine this possibility, we restricted the analysis to only inactive bout durations where flies exhibited movement in the 30 sec window prior to the 30 sec quiescent period before “laser on.” Normalized arousal thresholds of these similar duration inactive bouts exhibited a weak, but significant, negative correlation with ZT time (fig. S2C). Together, these data are consistent with multiple factors influencing arousal threshold: the light/dark transition (promoting arousal at ZT12), homeostatic sleep drive (gradually decreasing across the night), and the circadian clock (encouraging activity during the morning and evening peaks).

Prior work in *Drosophila* has largely used a “≥5 min inactivity” definition for sleep, where inactivity is assessed by whether a fly crosses a single IR beam (*24*, *50*). However, there have been suggestions that shorter time windows of inactivity may be associated with sleep in vinegar flies (*51*, *52*). Thus, using our IR laser-based closed loop system, we examined the duration of quiescence consistently associated with an elevated arousal threshold. We first labeled inactive bouts as “quiet wakefulness” (QW) if the fly did not exhibit locomotion during the 30 sec window prior to “laser on,” but performed microbehaviors including grooming, feeding, or defecation. The normalized arousal threshold for QW was low, as expected (Fig. 2E). We then assessed the duration of behavioral “quiescence” of individual perturbation bouts by determining if the fly exhibited grooming, feeding, defecation, or locomotion; a “quiescent” bout meant that the fly did not engage in any of those behaviors during the time window. Compared to QW, there was no difference in normalized arousal threshold if the flies were quiescent for 30-60s or 1-3 mins. In contrast, we found that flies that were quiescent for ≥3 min prior to laser on exhibited a marked increase in arousal threshold vs. QW (Fig. 2E). These data suggest that flies sleep in bouts of quiescence shorter than 5 min and argue that ≥3 min behavioral quiescence (defined as the absence of locomotion, grooming, feeding, or defecation) is associated with sleep in freely-moving *Drosophila*.

Previous studies have noted that flies tend to sleep near their food source (*53*). We examined the location of flies in our assay during inactive bouts lasting <1 min, between 1-3 mins, and >3 mins. Our data suggest that bouts of inactivity lasting <1 min are randomly dispersed throughout the chamber. In contrast, flies that were immobile for periods >1 min tended to stay near the food (Fig. 2F). Thus, these data suggest that flies prepare for sleep by engaging in pre-sleep quiescence behavior near food after 1 min inactivity, followed by sleep after 3 mins of quiescence.

Because our closed-loop IR laser-based system is precise and gentle and does not involve moving the fly, we were able to clearly visualize microbehaviors as the animal exited a sleep state, which has not been previously performed in *Drosophila*. After the laser turned on and gradually ramped up in intensity, we sometimes noted reversal of specific microbehaviors while the fly remained quiescent. For example, prior to the fly engaging in active locomotion, one can detect an “up”/dorsal movement of the antennae or halteres (Fig. 2, G and G’ and fig. S2, D-F). Interestingly, the haltere movement can occur on its own (fig. S2D) or precede the antennal movement (Fig. 2G’ and fig. S2E), prior to arousal. Alternatively, the haltere and antennal movements can occur simultaneously (fig. S2F). In contrast, during these transition periods, we never observed movement of the antennae before the haltere. These findings imply that HS are associated with a deeper stage of sleep and that subtle arousal stimuli can unmask different sleep sub-states.

We next asked whether reversibility of HS was specific for arousal from heat, or whether it could be seen with other arousing stimuli. We thus performed brief, repetitive (1 sec) mechanical stimulation every min using our imaging set-up. Interestingly, we observed quiescent animals with repeated haltere “up”/dorsal movements with the mechanical stimuli, followed by haltere down/ventral movements after the stimuli, suggestive of repeated microarousals (fig. S2, G and H). Taken together, our closed-loop system allows for precise quantification of arousal threshold and suggests the presence of different sleep sub-states marked by specific microbehaviors.

### Activation of dFB and R5 neurons leads to distinct microbehaviors

To demonstrate the utility of our high-resolution video imaging approach, we used FlyVISTA to characterize the behaviors observed following optogenetic activation of two proposed sleep-promoting circuits in *Drosophila*. In *Drosophila*, arguably the best-studied neural cluster suggested to promote sleep is a group of dorsal fan-shaped body (dFB) neurons. These neurons have been suggested to rapidly induce sleep-like behavior and improve memory performance in *Drosophila* (*36*, *54*, *55*). While a number of driver lines have been used to target these neurons to promote sleep, many groups have focused on *R23E10-Gal4*, as this line has relatively specific labeling of dFB neurons in the central brain (although connectomics data indicate that this driver line includes multiple distinct cell types) (*56*). Interestingly, recent papers have argued that the sleep-promoting effects of *R23E10-Gal4* are not attributable to the dFB neurons, but rather to neurons in the thoracic ganglion (*57*, *58*). To address the behavioral consequences of activating dFB (or the recently described subset of thoracic ganglion) neurons, we performed recordings consisting of 5 mins of baseline (“pre-stim”), 5 mins of 1 Hz optogenetic stimulation (“stim”), and 10 mins after stimulation (“post-stim”) for *R23E10-Gal4>UAS-CsChrimson* flies between ZT3-9. As a control, we also tested *empty-Gal4>UAS-CsChrimson* flies.

We first quantified the differences between these genotypes using centroid tracking data generated by our model. Optogenetic activation of both dFB and R5 neurons leads to an overall decrease in translational movement and total distance covered (Figs. S3, A-F), whereas no effect is seen on control animals. This reduced velocity persists following R5 activation, while in contrast velocity returns to near pre-stimulus levels for dFB activation. In addition, we asked if activation of these circuits influences their position in the arena, since flies sleep near food (Fig. 2F) (*53*). Interestingly, dFB activation leads to cessation of movement primarily in the bottom of the chamber or the sides further away from the food port, suggesting that flies avoid an upside-down orientation (Fig. S3G). In contrast, flies exhibit a more uniform distribution across the chamber following R5 activation, with some preferring to sleep near food without clear avoidance of upside-down orientation.

Next, we classified microbehaviors before, during, and after optogenetic stimulation. Strikingly, although optogenetic activation of *R23E10-Gal4>UAS-CsChrimson* flies led to a rapid cessation of gross locomotor activity (“moving”), it induced frequent micromovements (vertical movements of the thorax with jerky leg movements) and grooming, inconsistent with sleep behavior (Fig. 3, A and B and videos S3 and S4). These micromovements are distinct from leg adjustment behavior; micromovements involve the movement of the whole body along with the rapid movement of the legs (Fig. 3, C and D), whereas “leg adjustment” describes the brief movement of 1 or 2 legs during extended periods of quiescence. These observations are similar to findings in a previous report, suggesting thermogenetic activation of dFB neurons leads to increased micromovements (*59*). To quantify how the body of the animal moves during micromovement and leg adjustment behaviors, we trained a custom computer vision model, YOLO (You Only Look Once) (*60*) that performs object detection and segmentation. We used the segmentation masks to quantify how much of the body changes/moves, which revealed significantly higher total movements for micromovement than leg adjustment (Fig. 3E).

**Fig 3.**
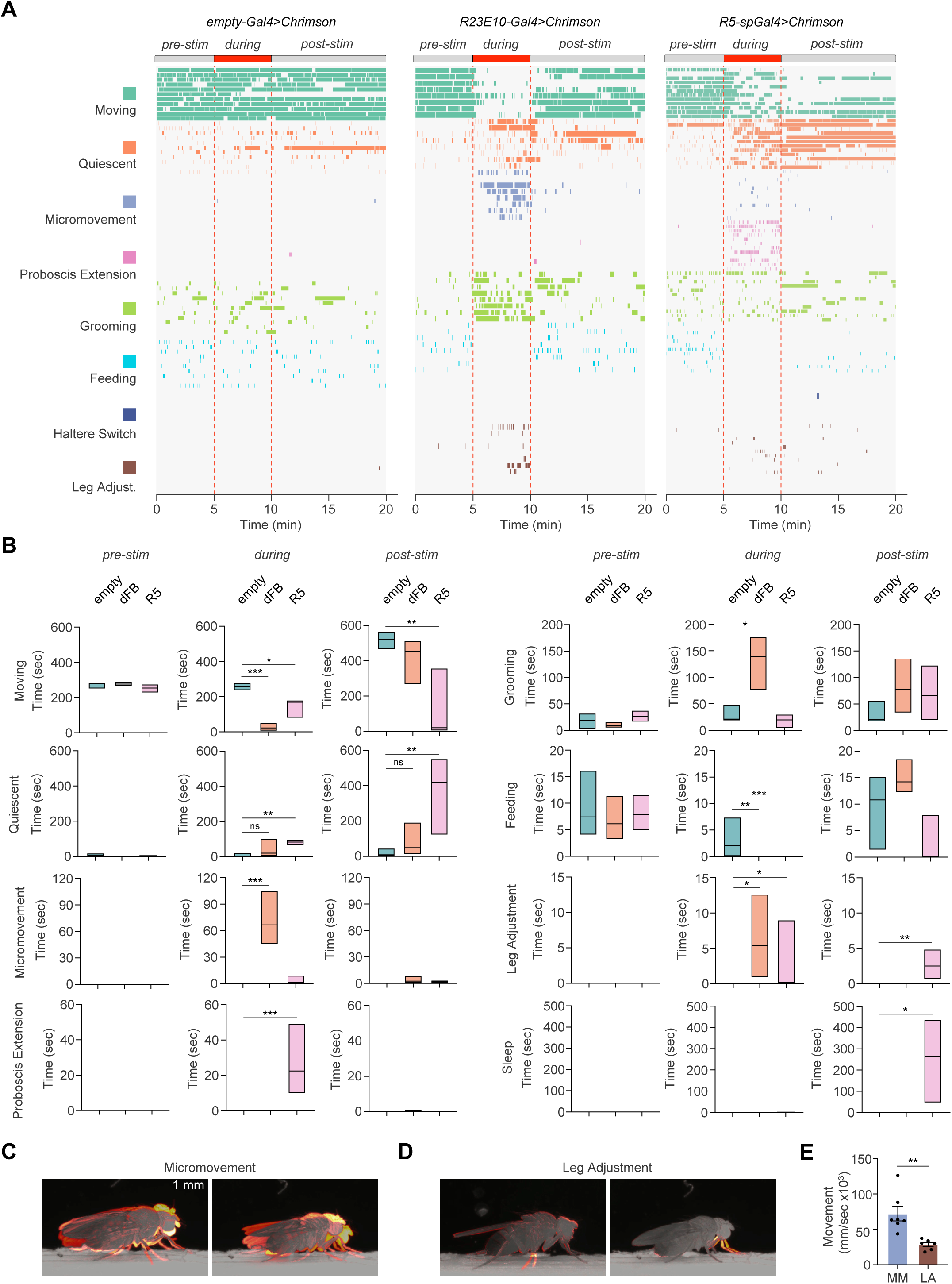
Optogenetic activation of putative sleep circuits promotes distinct microbehaviors. (**A**) Distributions of 8 distinct behaviors (moving, teal; quiescent, defined by absence of any observable movement, orange; micromovement, vertical thoracic movements with jerky leg movements, purple; PE, pink; grooming, green; feeding, crystal teal; HS, blue; and leg adjustment, brown) before, during, and after 5 min optogenetic stimulation (1 Hz) for control (empty-Gal4, n=11), dFB (*R23E10-Gal4,* n=8), and R5-splitGal4 (*R58H05-AD, R46C03-DBD*, n=12) male and female flies expressing CsChrimson at ZT3-9. (**B**) Duration of time spent in moving, quiescent, micromovement, PE, grooming, feeding, leg adjustment, and sleep states before (pre-stim), during (during), or after (post-stim) optogenetic stimulation for *empty-Gal4>UAS-CsChrimson* (empty, teal), *R23E10-Gal4>UAS-CsChrimson* (dFB, orange), and *R58H05-AD, R46C03-DBD* (R5, pink) flies. Kruskall-Wallis with post-hoc Dunn’s and Bonferroni correction. ns, not significant, *, *P*<0.05, **, *P*<0.01, and ***, *P*<0.001. **(C** and **D)** Motion heatmaps of 1 sec movement of the indicated behaviors. (**E**) Average movement of the tracked centroid per behavior (micromovement, purple; leg adjustment, brown). Unpaired t-test; **, *P*<0.01.

In addition, our group previously identified a set of ellipsoid body neurons (R5) proposed to be important for homeostatic sleep drive in *Drosophila* (*37*, *38*, *61*). Thus, we wished to use FlyVISTA to investigate the finer behavioral consequences of activating R5 neurons. Because it was recently found that some Gal4 drivers used to label R5 neurons were contaminated with arousal-promoting neurons, we used a split-Gal4 line (*R58H05-AD, R46C03-DBD*) that does not express in those additional cells (*37*, *62*). Our previous data suggested that thermogenetic activation of R5 neurons led to an increase in sleep during the heat treatment, followed by persistent sleep behavior after cessation of the heat treatment (*37*, *38*). 1 Hz optogenetic activation of *R58H05-AD, R46C03-DBD>UAS-CsChrimson* flies produced a significant reduction in locomotor activity during activation. Interestingly, these flies exhibited a robust increase in ∼0.3 Hz PE events during optogenetic stimulation (video S5). In contrast, no PE events were observed with *R23E10-Gal4* or *empty-Gal4* activation. Following stimulation of R5 neurons, there was a significant increase in persistent sleep (Fig. 3, A and B and video S6). Taken together, these data suggest that optogenetic activation using the *R23E10-Gal4* driver does not appear to promote sleep, while activation of R5 neurons leads to an interesting phenotype with PE events during stimulation, followed by persistent sleep after stimulation, consistent with a role for R5 neurons in sleep drive.

### Automated behavioral categorization using FlyVISTA

A central goal of computational ethology is the automated quantification of behavior in freely moving animals. Pose estimation tools, such as DeepLabCut (*8*) and SLEAP (*63*), have greatly facilitated the annotation of body parts from video data. However, it still remains a significant challenge to extract meaningful behavior from these annotations (*2*, *4*, *5*). This challenge is particularly pronounced when analyzing the microbehaviors associated with sleep. First, because we imaged the flies at high-resolution and over a long period of time, each dataset was large (∼10-20 GB for each 16 hr recording).

Second, the microbehaviors associated with sleep were small, sometimes involving movements as small as 40-50 microns. Third, because the animals were freely moving, the appearance of these movements was variable. Fourth, these movements occurred across a range of timescales, ranging from secs to mins.

Thus, we developed a novel computational pipeline for FlyVISTA, which classifies behaviors in unannotated fly videos using a set of manually annotated fly videos (which serve as a “gold standard”). Labels corresponding to one of five behavioral categories (PE, HS, leg adjustment, grooming, and feeding) are assigned to each microbehavior-containing frame in the video. We were unable to analyze postural relaxation (Fig. 1F) using this pipeline, because this movement is too subtle and occurs over a very long duration. The FlyVISTA pipeline involves three steps: a) feature extraction, b) semi-supervised behavioral embedding, and c) committee classification of behavioral categories (Fig. 4A). The pipeline takes 2D positional coordinates of the unannotated fly obtained from the DeepLabCut-generated pose estimations, and then computes various spatiotemporal features, such as distances between specific body-parts (fig. S4A). Wavelet transformations are applied on these raw features to extract spatiotemporal representations of each frame, resulting in a high-dimensional representation. To reduce the computational complexity and to increase predictive performance, we first classified frames into quiescent vs “active” (quiescent with microbehavior) frames and filtered out the quiescent ones lacking microbehaviors. In general, this classification was robust for most microbehaviors, but frames with HS movements were the most likely to be incorrectly classified as quiescent, given the chance for the halteres to be obscured by other body parts and the small movement involved (fig. S4B). Next, the high-dimensional representations were reduced to several lower-dimensional embeddings, where each unannotated fly video was paired with every annotated fly video, creating a comprehensive set of pairings. This pairwise strategy was adopted instead of a joint embedding strategy to prevent videos that are significantly dissimilar from distorting the embedding. Nearest neighbor classification was performed in this lower-dimensional space for each pair of annotated and unannotated videos, where each classifier presents a probability of categories for each frame of the unannotated fly. The final step involved combining the nearest neighbor classifications probabilistically within a committee of classifiers (Fig. 4A) to generate an “aggregated behavioral score” for each frame of the unannotated fly. A committee approach was used, because of the substantial variability of the behavioral repertoire in an individual fly. For example, analysis of the behavioral repertoire of an unannotated fly with HS was markedly hampered if compared to a single annotated fly exhibiting few or no HS during the night (fig. S4C).

**Fig. 4.**
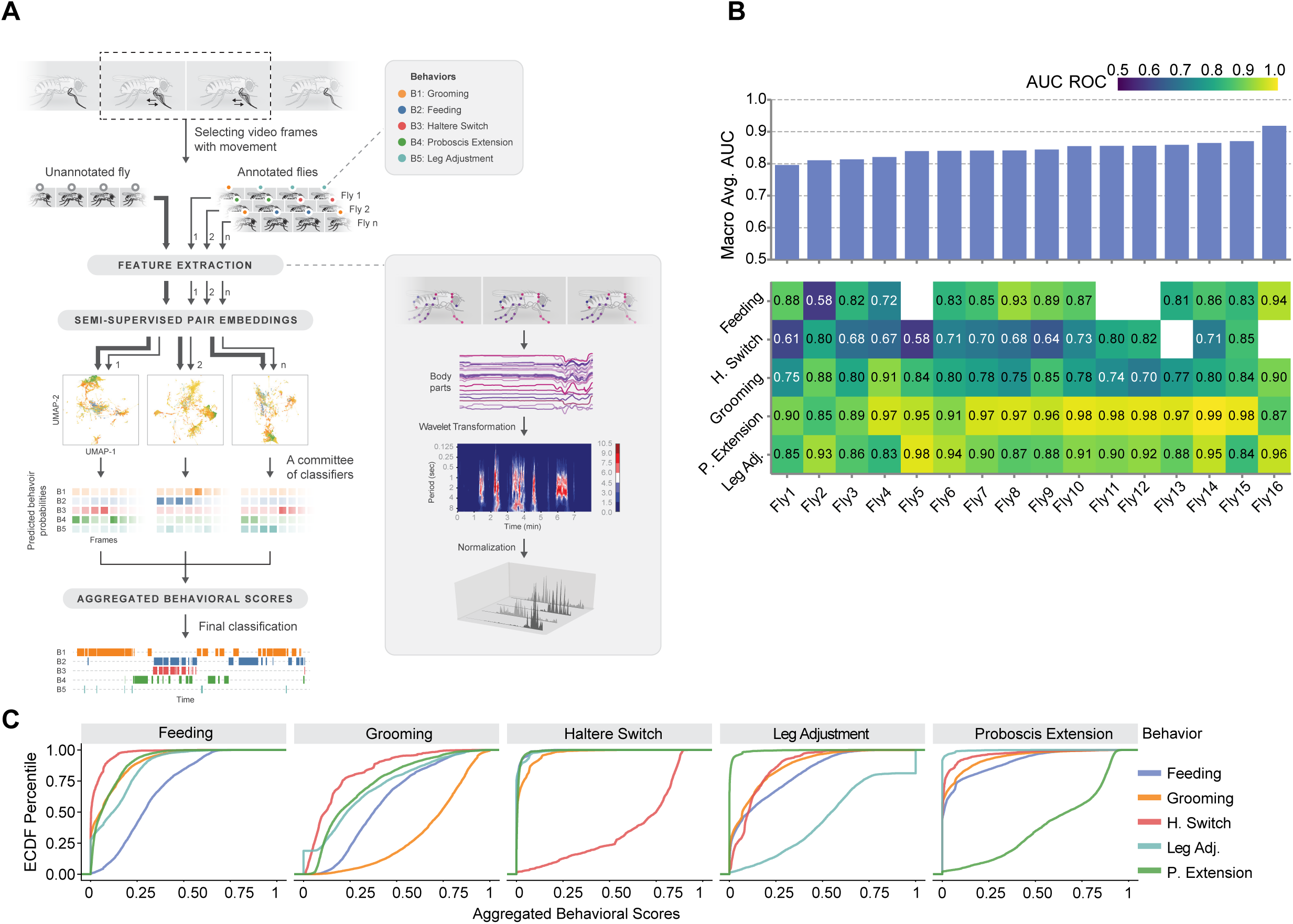
A semi-supervised computational pipeline to analyze sleep-associated microbehaviors in *Drosophila*. (**A**) Illustration of the main stages of the pipeline. To perform behavioral analysis, our pipeline starts by extracting meaningful spatiotemporal features from the body part positions, followed by a wavelet transformation and L1 normalization. Then, micro-activity detection is performed to distinguish quiescence and behaviors of interests using a random forest of decision trees. After that, semi-supervised embeddings of the time points detected as micro-activity are computed for each annotated and unannotated fly experiments separately. Finally, a committee of annotated fly experiments predicts behavior scores by performing a joint nearest neighbor analysis on the embedding spaces. The output of the pipeline is a distribution of scores for behavioral categories, per video frame. (**B**) Performance summary of behavior mapping with the area under curve (AUC) scores of receiver operating characteristic (ROC) for 16 experiments. (Below) Each column of the heatmap corresponds to a leave-one-out experiment, and each value measures the AUC of ROC curves for different behavior categories. (Above) bar-plots aggregate AUC values as a macro average. Absent behavior categories are left blank for some experiments. (**C**) Empirical cumulative distribution function plots of aggregated behavioral scores of each behavioral category (PE, green; HS, red; leg adjustment, teal; feeding, blue; and grooming, orange) in all leave-one-out experiments combined. Each plot demonstrates the aggregated behavior score distributions of all the time points with the corresponding true annotation. These distributions reveal the predictive power of the scores, especially for PE, HS, and grooming.

To assess the predictive performance of behavior classification by FlyVISTA, we performed a leave-one-out cross-validation on the 16 annotated fly videos (Fig. 4B). Specifically, we used 15 manually annotated videos to predict the behavior of the held-out fly video. The FlyVISTA pipeline demonstrates strong performance in classifying PE, achieving AUC ≥ 0.95 for 11 splits out of 16 splits. Similarly, for leg adjustment, the AUC exceeds 0.9 for 9 splits and only drops below 0.85 in one split. The AUC score for grooming detection ranges from 0.70 to 0.91, performing slightly worse than for PE and leg adjustment. For grooming, behavioral scores incorrectly predicting feeding and, to a lesser extent PE and leg adjustment, were relatively high, which results in false positives leading to the decrease in performance (Fig. 4C). The AUC scores for feeding are above 0.7, except for one split out of 13 in which feeding was observed. The most challenging behavior for FlyVISTA is HS, as it is the most nuanced behavior. The AUC score for HS never exceeds 0.85, but is above 0.7 in 8 splits out of 14 splits. Overall, the macro average score is above 0.8 for 15 splits. We also calculated precision-recall and receiver operating characteristic (ROC) curves for each split, as well as the interpolated weighted averages of the curves, using the generated behavioral scores (fig. S4D). The maximum F1 score for grooming was 0.6 and did not exceed 0.4 for HS. In contrast, PE and leg adjustment were most robustly detected, achieving F1 scores >0.8 and 0.85, respectively for some splits. Overall, these findings argue that FlyVISTA robustly classifies wake- and sleep-associated microbehaviors in *Drosophila*.

### Quantification of fly sleep using FlyVISTA

We next used FlyVISTA to quantify sleep in freely moving flies (using our definition of ≥3 min quiescence with no feeding or grooming). First, we examined the organization of sleep/wake in these animals. Compared to sleep observed in fixed, tethered flies (which is highly fragmented) (*35*, *52*, *64*), we found that sleep in freely moving flies was relatively consolidated (Fig. 5A). We plotted sleep amount and bout durations across ZT time (Fig. 5, B to E). Female flies sleep at a consistently high level throughout the night, while male flies exhibit a sharp reduction in sleep amount in the late night (Fig. 5C), consistent with pronounced morning anticipation seen in males (*45*, *65*). We next compared baseline sleep features measured in freely moving flies with FlyVISTA vs using traditional analyses using glass tubes in *Drosophila* Activity Monitors (DAM, Trikinetics). Baseline sleep amount at night was significantly higher using the Trikinetics system in both males and females (fig. S5, A, B, E, and F). Moreover, sleep consolidation, as assessed by characterizing sleep bouts, was markedly overestimated using Trikinetics monitors (fig. S5, C, D, G, and H). Interestingly, the range of sleep bout duration measured using FlyVISTA was narrow, suggesting that flies have a typical sleep bout length (∼19 mins and ∼27 mins for females and males, respectively). One of the key defining features of sleep is its regulation by homeostatic forces (*66*). Thus, we also assessed sleep amount and consolidation following mechanical sleep deprivation (SD) from ZT12-ZT24. We calculated sleep amount and bout duration from ZT0-ZT2 in the presence or absence of SD. As expected, a significant increase in both sleep amount and bout duration was observed in both males and females following SD (Fig. 5, F-H). Together, these findings suggest that fly sleep measured using FlyVISTA is more natural and consolidated compared to when flies are tethered, but more fragmented with consistent bout durations compared to Trikinetics measurements.

**Fig. 5.**
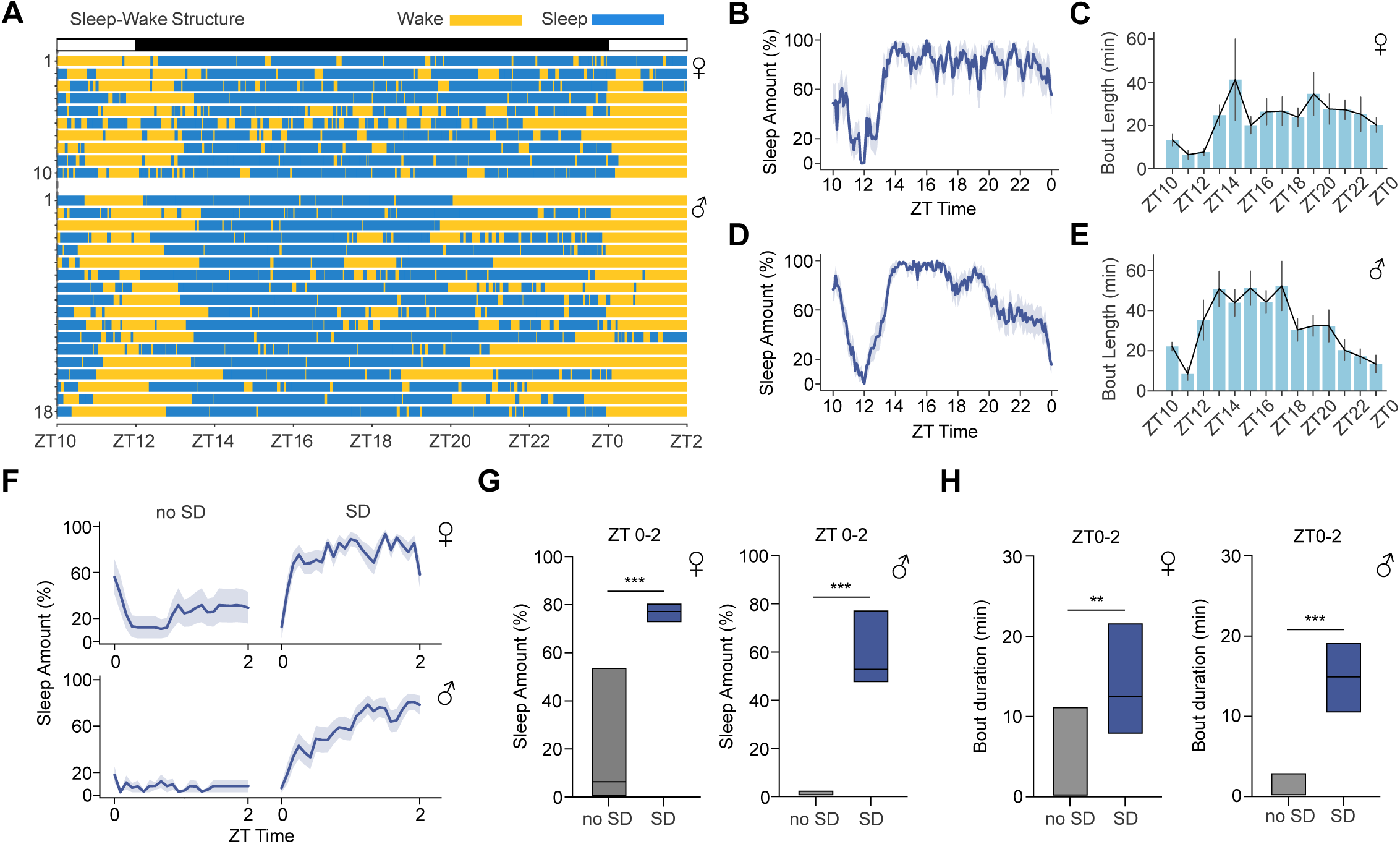
Quantification of sleep behavior using FlyVISTA. (**A**) Distribution of wake (yellow) and sleep (blue) bouts plotted for female and male flies for the ZT times shown. (**B** and **C**) % sleep in 5 min bins (**B**) and sleep bout duration in 30 min bins (**C**) from ZT10 to ZT0 for female (n=10) flies. Shading and error denote SEM. (**D** and **E**) % sleep in 5 min bins (**D**) and sleep bout duration in 30 min bins (**E**) from ZT10 to ZT0 for male (n=18) flies. Shading and error denote SEM. (**F**) % sleep from ZT0 to ZT2 in 5 min bins in the presence (SD) or absence (no SD) of 12 hr SD from ZT12-ZT24 for female (top, n=10 for no SD and 17 for SD) and male (bottom, n=18 for no SD and 19 for SD) flies. Shading denotes SEM. (**G** and **H**) Simplified box plots showing % sleep amount (**G**) or sleep bout duration (**H**) from ZT0 to ZT2 in the presence (SD) or absence (no SD) of 12 hr SD from ZT12-ZT24 for the female (left) and male (right) flies in (**A**). Simplified box plots denote 75^th^, median, and 25^th^ percentiles. Mann-Whitney U tests; **, *P* < 0.01 and ***, *P* < 0.001.

### Proboscis extension behavior is under provoked, but not baseline, homeostatic control

A prior study using tethered flies argued that sleep associated-PEs decrease monotonically during the night under baseline conditions and are increased following SD (*35*). To characterize PE microbehaviors in freely moving flies, we utilized FlyVISTA to quantify these behaviors under baseline conditions and after SD. In male and female flies under baseline conditions, we found that PE frequency peaked at ∼17 events/hr (females) or ∼30 events/hr (males) and did not demonstrate a monotonic decrease across the night (Fig. 6, A and B). Instead, interestingly, there appeared to be peaks of PE events in the early and middle portions of the night (Fig. 6B). The mid-night peak occurs during a period of low locomotor activity (Fig. 1D), suggesting that it is not simply linked to physical exertion.

**Fig. 6.**
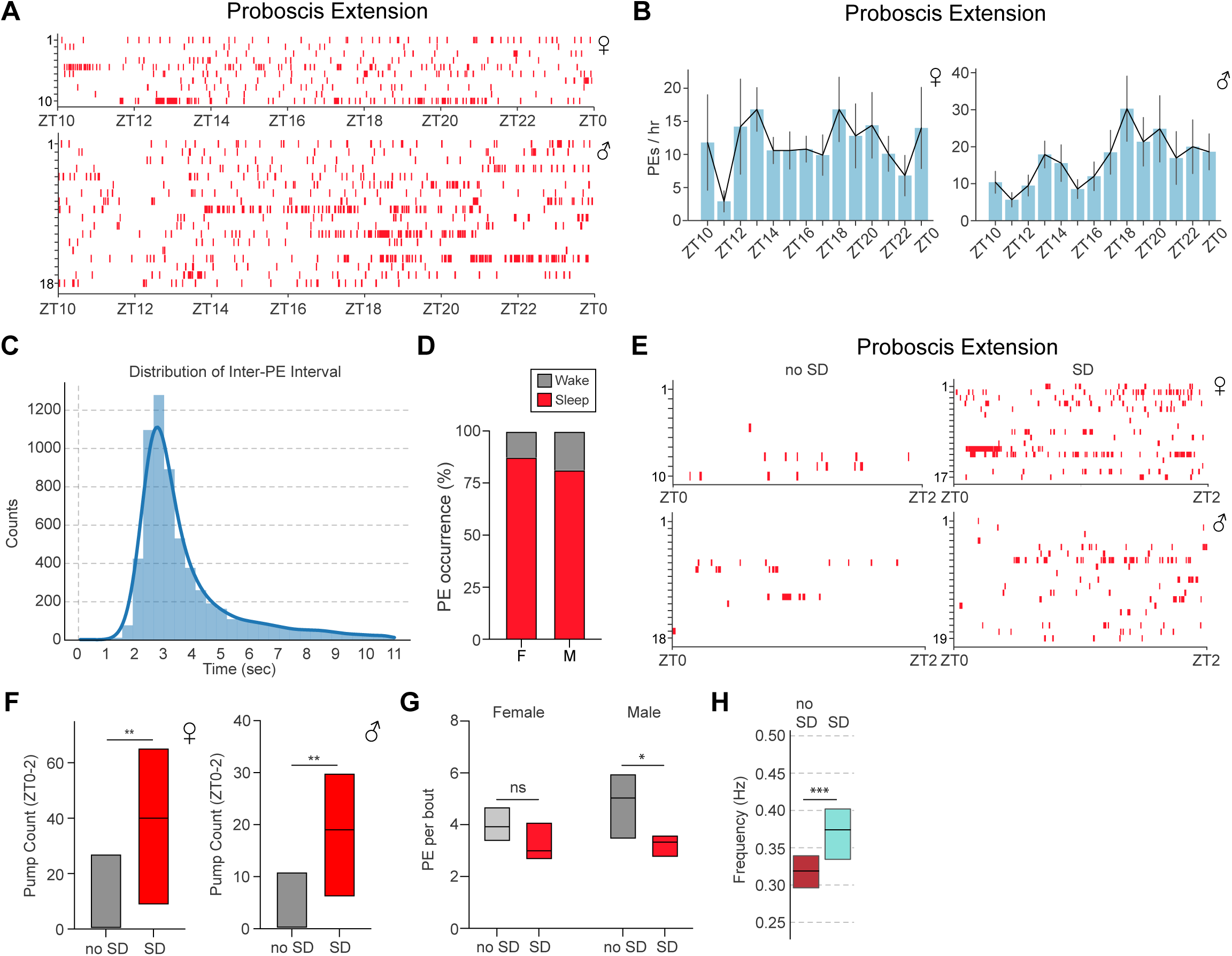
Characterization of proboscis extension behavior. (**A**) Distribution of PE events across the night. Individual PE events (red) plotted for each individual female (upper panel) and male (lower panel) fly from ZT10 to ZT0. Data are from the same flies as in Fig. 5A-5E. (**B**) PE/hr from ZT10 to ZT0 for the female (left) and male (right) flies in (**A**). Error denotes SEM. (**C**) Histogram showing distribution of inter-PE intervals, with a peak near 3 sec. (**D**) Stacked bar plot showing proportion of PE events occurring during wakefulness (gray) or sleep (red) for female (F) and male (M) flies. (**E**) Distribution of PE events (red) from ZT0-ZT2 in the presence (SD) or absence of 12 hr SD (no SD). Individual PE events plotted for each individual female (upper panel) and male (lower panel). Data are from the same flies as in Fig. 5F-5H. (**F**) Simplified box plots showing PE count from ZT0 to ZT2 in the presence (SD) or absence (no SD) of 12 hr SD for the female (left) and male (right) flies in (**E**). Mann Whitney U-test; **, *P* < 0.01. (**G**) Simplified box plots showing the number of PEs per bout in the presence (SD, red) or absence (no SD, gray) of 12 hr SD from ZT12-24 for female (n=10 and 17 for no SD and SD) and male flies (n=18 and 19 for no SD and SD). Mann Whitney U-test; ns, not significant and *, *P*<0.05. (**H**) Simplified box plot showing frequency of PE per bout in the absence (gray, no SD, ZT10-ZT2, n=28) vs presence (red, SD, ZT0-ZT6, n=36) of 12 hr SD from ZT12-ZT24. Data are pooled data from males and females. Top and bottom of the boxes represent 75^th^ and 25^th^ percentiles, and middle line represents median. Mann Whitney U test; ***, *P* < 0.001.

Previous work showed that these PEs typically occur with ∼3s inter-PE interval (IPI) (*35*, *47*), and similar results were obtained using FlyVISTA (Fig. 6, C and H). There is some variance in the IPI, which can occur within a single bout (fig. S6A), but also consistently between the first 2 PEs and the last 2 PEs, with the last IPI being significantly longer (fig. S6, B and C). PEs are not exclusive to sleep and can also be observed during wakefulness or after prolonged flight (*35*, *46*, *47*). Thus, we asked what proportion of PEs occurred during sleep vs wakefulness. We found that the majority (∼82% and ∼88% for males and females, respectively) occurred during sleep bouts (Fig. 6D). Because PE bouts are typically brief (∼10s), we had only 5 bouts where the laser turned on while the fly was exhibiting PEs. The mean normalized arousal threshold was higher than that observed for nearby QW bouts, but this difference was not significant (0.70 ± 0.36, n=5 sleep bouts with PEs vs 0.20 ± 0.36, n=9 adjacent QW bouts without PEs, *P*=0.38).

To investigate how these PEs were affected by SD, we performed mechanical SD from ZT12-ZT24 and characterized PE behavior using FlyVISTA. As has been previously described (*35*), SD triggered a substantial increase in PE count in both female and male flies, which persisted for at least 2 hrs after cessation of the SD (Fig. 6, E and F). Interestingly, SD also decreased the number of PEs per bout in males and led to a significant shortening of the IPI (Fig. 6, G and H). Taken together, these data reveal that PEs usually occur during sleep and are regulated by strong, but not baseline, homeostatic drive.

### Haltere Switch Behavior Defines a Deeper Sleep Stage in *Drosophila*

Finally, we characterized HS behavior using FlyVISTA. HS events also appeared to display a peak during mid-night in both males and females (Fig. 7, A and B). Haltere “up” events were identified less frequently than haltere “down” events (Fig. 7A), due to the difficulty identifying those events in a fly that has awakened and started walking. In addition, the FlyVISTA pipeline was trained to detect fast HS movements, and some HS “up” events are gradual and slow (fig. S7A). Interestingly, HS events were less commonly observed in males, which likely reflects a combination of fewer HS seen in this sex, as well as the challenge in visualizing haltere movements (males have more compact bodies, and halteres are more easily hidden behind legs or can rest on the posterior legs). The frequency of HS events was increased following SD in male flies, and there was a trend for an increase in HS events following SD for female flies (Fig. 7, C and D). Together, these data suggest HS microbehaviors are likely under homeostatic control.

**Fig. 7.**
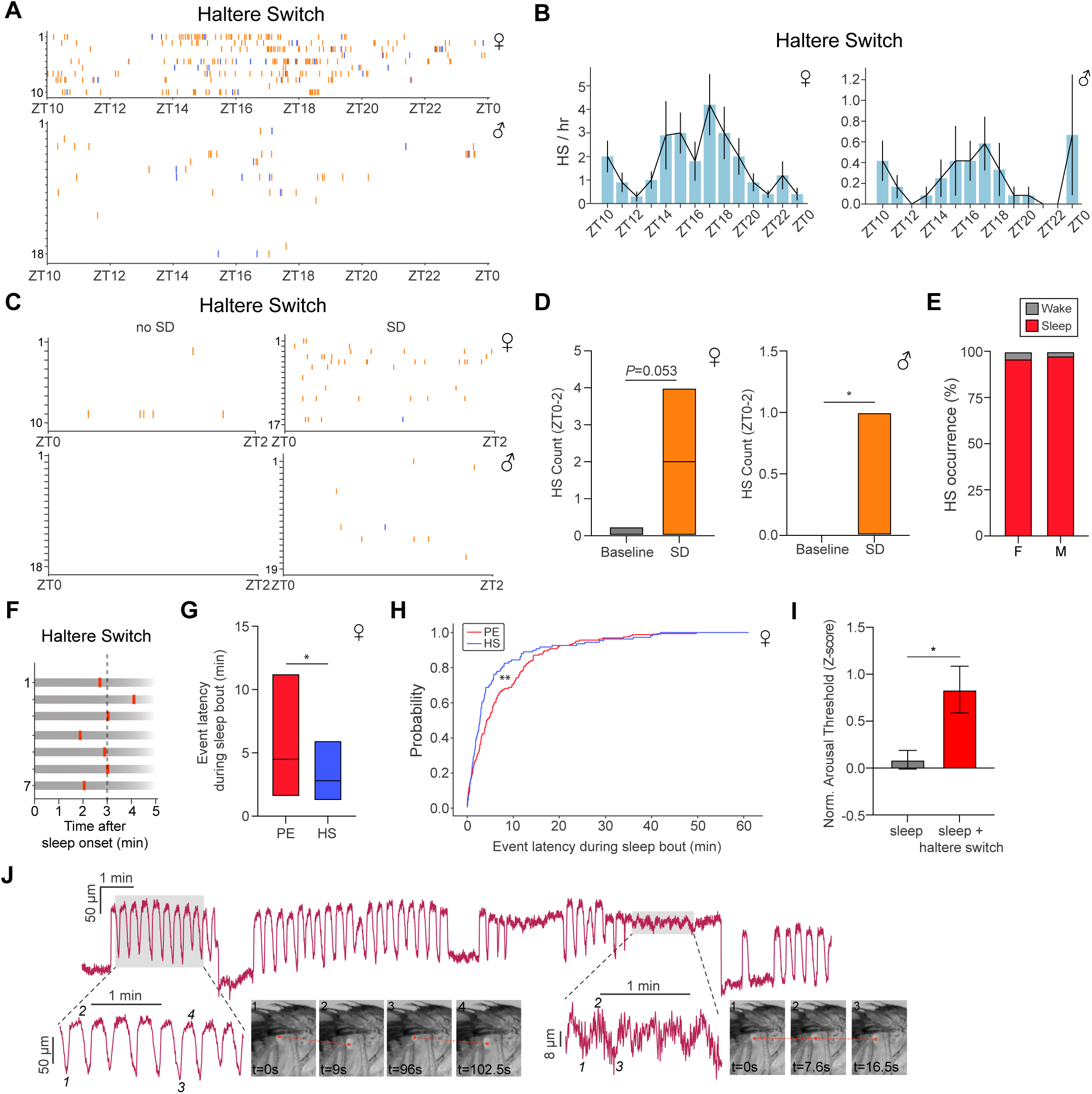
Haltere switch behavior identifies a deeper sleep state in *Drosophila*. (**A**) Distribution of HS events across the night. Individual HS plotted (orange ticks for “haltere down/ventral” and blue ticks for “haltere up/dorsal”) for each individual female (upper panel, red) and male (lower panel, blue) fly from ZT10 to ZT0. Data are from the same flies as in Figs. 5A-5E. (**B**) HS/hr from ZT10 to ZT0 for the female (left) and male (right) flies in (**A**). Only HS “down” events are plotted. Error denotes SEM. (**C**) Distribution of HS events from ZT0-ZT2 in the presence (SD) or absence of 12 hr SD. Individual HS events (orange ticks for “haltere down/ventral” and blue ticks for “haltere up/dorsal”) plotted for each individual female (upper panel) and male (lower panel). Data are from the same flies as in Figs. 5F-5H. (**D**) Simplified box plots showing HS count from ZT0 to ZT2 in the presence (SD) or absence (no SD) of 12 hr SD for the female (left) and male (right) flies in (**C**). Simplified box plots denote 75^th^, median, and 25^th^ percentiles. Mann Whitney U-test; *, *P* < 0.05. (**E**) Stacked bar plot showing proportion of HS (down) events occurring during wakefulness (gray) or sleep (red) for female (F) and male (M) flies. (**F**) Representative examples of HS timing relative to the onset of a sleep bout. (**G**) Simplified box plot (75^th^ percentile, median, 25^th^ percentile) showing latency of PE or HS events relative to the onset of a sleep bout in female flies. Mann Whitney U-test; *, *P*<0.05. (**H**) Cumulative distribution function plot for PE (red) and HS (blue) probability vs latency within a sleep bout. Kolmogorov-Smirnov test; **, *P*<0.01. Data in (**F**-**H**) are from the same flies as in Figs. 5A-5E. (**I**) Normalized arousal threshold for female flies identified as sleeping (≥3 min quiescence starting from “laser on”) in the absence (n=93) or presence (n=11) of HS behavior. Error denotes SEM, unpaired t-test; *, *P* < 0.05. (**J**) Haltere oscillations of different amplitudes across a sleep bout. Time series data show distance between tracked points in the haltere and scutellum. Shaded areas represent the enlarged time traces (below) with the accompanying representative images on the right. Smaller amplitude oscillations are shown on the right with the tracked haltere points denoted on the representative images.

Notably, in contrast to PEs, HS (haltere “down/ventral”) events are tightly associated with sleep (∼98% in males and ∼93% in females) (Fig. 7E) and exclusively seen during quiescence. We next examined when HS events occur following sleep onset. HS events are relatively early events within a given sleep bout, with the first HS event occurring ∼3 mins after sleep onset (Fig. 7, F and G). In contrast, PEs typically occurred later within a sleep bout, with the first PE in a sleep bout ∼5 mins after sleep onset. Indeed, the latency to PE was greater than the latency to HS during a sleep bout (Fig. 7, G and H). We next asked whether HS affected the depth of sleep (as defined by changes in arousal threshold). To do this, we identified sleep bouts and then compared arousal threshold levels if HS was present at “laser on.” We found that when flies were asleep and their halteres were “down,” they demonstrated an elevated arousal threshold level, compared to sleep bouts where halteres were “up” (Fig. 7I); these data suggest that HS “down” marks a deeper sleep stage.

Closer inspection revealed different subtypes of HS events during behavioral quiescence (fig. S7A). In addition to unitary HS events, flies exhibited spontaneous events where halteres briefly moved “up”/dorsal and then “down”/ventral; these episodes likely represent microarousal events, similar to those provoked by arousing stimuli (Fig. 2G and fig. S2, D-H). Conversely, sometimes following a HS “down” event, halteres would slowly move “up”/dorsal. Other times, multiple HS events would spontaneously occur in succession (“active haltere”) in a non-rhythmic manner (fig. S7B), as if the fly were repeatedly entering and exiting different sleep states (reminiscent of the repeated HS events with multiple arousal events; fig. S2, G and H).

Finally, distinct “haltere oscillation” events were also observed, where the halteres would slowly and rhythmically move up and down. The frequency of these HS oscillations was ∼0.09 Hz and could vary between ∼0.06 and 0.14 Hz within an individual animal and across time (fig. S7, C and D). As we examined these events more closely, we noted that these oscillatory haltere movements were usually, if not always, accompanied by rhythmic movements of multiple body parts (e.g., head, antenna, thorax, abdomen) (fig. S7, E and F). In unpublished data from an ongoing study, we have noted tonic activation of scutellar pulsatile organs (which function as “wing hearts” to pump hemolymph into the wings) (*67*) that lasts 6-7 sec, followed by 6-7 sec of complete inhibition of these muscles. This rhythmic activity matches the frequency of the “haltere oscillations” that we observe, and so we hypothesize that this slow oscillatory rhythm reflects the gradual filling and emptying of the scutellar hemocoel. Intriguingly, during sleep, flies sometimes exhibited periods where these ∼0.09 Hz haltere oscillations were several-fold greater in amplitude (Fig. 7J), suggestive of periods of sleep where the fly musculature may be highly relaxed. Taken together, these data reveal that HS events marks a deeper sleep stage in flies and suggest that visualizing haltere dynamics may uncover novel sleep sub-states in *Drosophila*.

## Discussion

> *"The horse will sleep standing in the warm shelter of the stable, though it lies down in the pasture; the bird reposes perching, but with its head buried in the feathers of the wing; the serpent coils itself in a circle, or folds itself into the smallest possible space….. The insect withdraws from the scenes of its ordinary activity, and is in a state of somnolent rest when it remains motionless. As the insect has no eyelids, no external closure of the eye gives evidence of sleep."*

> Richard Hill, Annual Of Scientific Discovery, 1864

Sleep has been reported to occur in widely divergent animal species (*13–17*), and the challenge of visually identifying sleep in insects has been long recognized. From these many species, the most commonly utilized non-mammalian model organism to study sleep is *Drosophila melanogaster* (with > ∼1,200 citations in Pubmed in 2024) (*24*, *25*, *50*). The vinegar fly system combines a powerful genetic toolkit with a compact nervous system and a recently defined connectome (*56*, *68*), making it an attractive system to investigate the function and regulation of sleep. However, over the past two decades, the vast majority of *Drosophila* sleep studies have utilized a simple operational definition, which claims that sleep is defined by a ≥5 min period of behavioral quiescence, associated with an increase in arousal threshold (*24*). The ease and simplicity of this approach has enabled large-scale genetic and circuit screens (*38*, *69*– *74*). However, the uniform application of this 5 min definition has likely led to sleep in *Drosophila* not being robustly characterized, limiting the potential of the fly model to understand the nature and function of sleep.

Here, we apply a computational ethology-based approach to study sleep in *Drosophila*. While video-based machine-learning algorithms are increasingly being used to quantify the structure of behaviors in freely moving animals (*4*, *75–77*), the study of sleep using this approach poses a particularly difficult problem; outside of a few exceptions (*78*, *79*), sleep is a quiescent behavior and so annotation and classification has to be performed on “microbehaviors” (small subtle behaviors occurring during sleep).

These sleep-related microbehaviors could provide insights into the different sub-states occurring during fly sleep. Moreover, unlike many “active” behaviors (e.g., locomotion, grooming, courtship, feeding, aggression) that are more temporally discrete, sleep behaviors occur over a longer timescale. To address these computational challenges, we developed FlyVISTA, which integrates a high-resolution closed-loop video imaging system with a pose estimation model and a novel semi-supervised computational pipeline to extract microbehaviors occurring during sleep.

We used FlyVISTA to perform detailed characterization of the effects of circuit manipulations on sleep in flies. Strikingly, we find that optogenetic activation using the *R23E10-Gal4* driver--one of the most commonly used drivers to promote fly sleep— reduces locomotion, but does not produce behavioral quiescence; instead, frequent micromovements and grooming behavior are observed. This driver has been used to label the dFB neurons proposed to promote sleep in flies, but recent data have argued that specific thoracic ganglion neurons (not the dFB) neurons are actually the neurons that induce sleep (*57*, *58*). Regardless of the neurons implicated, our data go beyond these recent assertions and suggest that the activating neurons labeled by the *R23E10-Gal4* driver does not induce sleep. In contrast, optogenetic activation of R5 neurons reduces locomotion and triggers PEs during activation, followed by persistent sleep. These data are consistent with the notion that R5 activation induces sleep need, as PEs are increased following SD (*35*) (Fig. 5) and have been suggested to facilitate waste clearance related to sleep (*35*). The subsequent persistent sleep behavior could thus be driven by this increase in sleep need. How these R5 neurons, which get strong visual input similar to other ring neurons, integrate visual cues, produce PEs, and coordinate sleep drive remains an interesting question to resolve (*81–83*).

Using FlyVISTA, we made several interesting observations regarding the nature of fly sleep and its associated microbehaviors. First, we found that flies can sleep in relatively short bouts (≥3 mins) of quiescence. This finding is consistent with recent work suggesting that flies can sleep in bouts less than 5 min (*51*, *52*). Together, these data suggest that fly sleep is underestimated using the widely-used 5 min definition.

Second, we confirmed the presence of PE-associated sleep occurring in freely moving flies. However, previous work using a tethered preparation suggested that flies exhibited frequent PE (up to ∼100 PE events/hr) (*35*). In contrast, we find several-fold fewer PE events in freely moving flies. PEs can be observed during wakefulness and be triggered by metabolically-demanding activities, such as flight (*47*). Since tethering a fly is likely stressful and may increase metabolic demand due to increased locomotion, some of the PE events seen in tethered flies may be unrelated to sleep *per se*. Third, FlyVISTA enables examination of the spatiotemporal structure of fly microbehaviors. For example, we distinguish multiple oscillatory body movements during fly sleep. Fast rhythmic movements of the legs and slow whole-body oscillations reflect actions of various pulsatile organs. Of note, the slow whole-body oscillations likely underlie previously described periodic antennal movements seen during fly sleep (*64*). Haltere movements during quiescence also comprise a heterogenous group of behaviors. Halteres can undergo “up” and “down” movements with different kinetics, suggestive of microarousals or microsleeps. Intriguingly, there are also periods of behavioral quiescence where the halteres and other body parts exhibit large oscillatory movements, suggesting that the body of the fly is particularly relaxed during these periods (Fig. 7J). It is tempting to speculate that these intervals reflect sleep stages with lower muscle tone.

Importantly, we define a novel microbehavior (HS) which, unlike PE, is exclusively seen during quiescence. Sleep associated with HS exhibits a greater arousal threshold than sleep without HS, suggesting that the HS “down” microbehavior defines a deeper stage of sleep. Interestingly, we also find that the first HS event in a given sleep bout occurs ∼3 mins after sleep onset, suggesting that entrance into this deeper sleep stage occurs relatively early within a sleep bout. These findings, coupled with prior work proposing similar findings with PE-associated sleep, suggest that multiple sleep stages exist in *Drosophila*. Indeed, a classic study in humans > 150 yrs ago predicted the cycling of sleep stages by simply assessing arousal thresholds in response to varying sounds from a hammer striking a slab (*83*). Other studies performing arousal perturbations and brain-wide Ca^2+^ imaging have also suggested the possibility that fly sleep has multiple stages (*51*, *52*). However, ultimately, fully characterizing different sleep stages in *Drosophila* will require neurophysiological imaging of both brain and muscle, preferably in freely moving animals.

Although we developed FlyVISTA to characterize fly sleep, there should be broad utility of this system for *Drosophila* neuroscientists. Among video imaging methods in *Drosophila*, FlyVISTA is unique in its use of a side-view chamber. Analyzing side-view images is more technically challenging (due to the increased likelihood of occlusion of body parts), but can provide a richer behavioral repertoire, such as HS events which cannot be visualized from top-view. Thus, FlyVISTA can be used to quantify, not just sleep, but potentially any behavior exhibited by freely moving flies, and our high-resolution video dataset (∼100 million frames) can be mined for additional behaviors that have not been previously described. In addition, while pose estimation-based video analyses of freely moving animals are being increasingly used, FlyVISTA is the first such method capable of characterizing microbehaviors in largely quiescent animals. Because important internal states can be revealed through subtle changes in posture and position of body parts, further development of applications like FlyVISTA should yield novel insights into how animals integrate internal states and external perception to generate behavior.

## Materials and Methods

### Fly strains

Flies were maintained on standard food containing molasses, cornmeal, and yeast at room temperature. *iso^31^* was used as the background strain (*84*), and all strains used were backcrossed into the *iso^31^* background at least 4 times. Unless otherwise specified, all flies used were 4-7 days old. *R23E10-Gal4* (BL #49032), and *empty-Gal4* (BL #68384) were obtained from the Bloomington *Drosophila* stock center. The *R58H05-AD, R46C03-DBD* driver was previously described (*37*). *UAS-CsChrimson-mCherry (VK5)* was a gift from Vivek Jayaraman.

### Behavioral Chambers

Flies were raised at 23 C° in 12:12 LD cycles in an environmental incubator. For each experiment, the corresponding fly was placed in a custom designed 3D-printed (Ultimaker 3) chamber with a 2.8mm (D) x 4.9mm (H) x 7.1mm (W) at the front. The chamber tapered so that at the back width was 4.1mm and height is 6.6 mm. A laser cut square acrylic window (7×7mm with a 2 mm thickness) was inserted at the front and the back of the chamber to keep the fly enclosed while allowing for recordings of behavioral videos. In some cases, these windows were coated with Sigmacote (SL2, Sigma Aldrich) to prevent flies from hanging onto the windows.

Individual 5-7 day old male and mated female flies were loaded into individual chambers 1-3 hrs prior to recording start time using either a mouth pipette or a small vacuum pump (FV-10, Furoro). A small capillary containing <5 ul of 2.5% yeast and 2.5% sucrose liquid food was inserted through a food port located at the top of the chamber. This capillary was secured using a plastic pipette tip or UV curable glue (3972, Loctite) to the chamber. Chambers were then attached to a 3D printed platform secured on an aluminum breadboard using 1/4" x 1/8" x 1/16" magnets. A telecentric lens (#63-074, Edmund Optics) attached to a machine vision camera (BFS-U3-23S3M-C, FLIR) was used to record 16 hrs (ZT10-ZT2) of behavior in each fly. An 850 nm ring light (LDR2-70IR2-850, CCS) was used to illuminate the fly. Camera settings, start/stop time and video encoding settings are controlled either through SpinView (FLIR) or custom-written MATLAB and Python scripts. All videos were recorded at 30 Hz.

### FlyVISTA for Behavioral Identification

#### Pose Estimation

We used DeepLabCut (*8*) for pose estimation. Of the 1304 labeled images from 88 files, 5% were held out for testing and the remaining images were used for training. We trained a ResNet-50 based neural network using batch size 4 and 200000 iterations. The resulting network had a train error of 6.31 pixels and a test error of 9.96 pixels for image-size of 1100×800 pixels.

#### Feature Extraction

The first stage of the FlyVISTA pipeline is feature extraction (*85*– *87*). This step comprises four consecutive steps: preprocessing, spatiotemporal feature computation, wavelet transformation, and normalization.

#### Preprocessing

The preprocessing step addresses two challenges: the occluded body-part problem and the noisy nature of pose estimation data. We utilized the confidence scores provided by body-part tracking software DeepLabCut (*8*) to detect which one of the left & right counterparts of a body part was occluded at a specific time point.

To filter data, we first detected erroneous “jumps” in tracking by comparing the estimated pose values to the median pose value of the same body part within a 0.5 sec window. If the pose estimation at a given time point exceeded the median value of the window centered at that time point by 15 microns, we considered it to be a “jump”, and removed the corresponding data point. The removed time points, i.e., jumps and occlusions, were imputed by interpolation. Following imputation, we applied a median filter (of size 0.2 sec) followed by a boxcar filter (of size 0.2 sec) to eliminate the rapidly changing signals without smoothing out short-duration low-amplitude behavioral signals, e.g., HS.

#### Spatiotemporal feature computation

In the subsequent step, we computed 13 meaningful and representative spatiotemporal features from the preprocessed tracking data. These features encompass measurements like the separation between the thorax and haltere, or the distance from the head to the proboscis (fig. S3A). To refine our behavioral representation, we applied wavelet transformation to capture postural dynamics over multiple timescales simultaneously (*85–87*). We applied continuous wavelet transformation with Morlet wavelet, that span 20 different frequency channels dyadically spaced between 1 Hz and 20 Hz. This procedure yielded a total of 260 feature values (13 features × 20 frequency channels). We normalized the power spectrum of different timescales as described before (*88*) and finally, we applied L1 normalization to generate a vector wherein the values were summed to 1 for each time point.

#### Elimination of quiescence frames that lack micro-activity

We first eliminated frames of quiescence devoid of micro-activity. This elimination served two purposes. First, given the high frame rate and long duration of video recordings, computational feasibility was a key concern. As more than 90% of the frames were either predominantly quiescent or contain macro-activities such as walking, filtering quiescence frames reduced computational demands significantly. Second, since quiescence frames mostly encompass noise (i.e., no relevant signal that relates to micro-activity), if they are included in normalization the amplified noise generates a uniform-like probability distribution for behavioral representation as was observed previously (*87*). This critically impacts the subsequent behavioral classification. Therefore, eliminating frames of pure quiescence devoid of micro-activity aims to circumvent this issue. To do so, we trained a random forest classifier (*89*), with 10 estimators (i.e., trees) with a maximum depth of 5, with Gini index as the impurity criterion, and classified quiescence vs activity frames.

#### Behavioral classification

Following the identification of frames with micro-activities, we performed dimension reduction on the high dimensional feature space from the power spectra of multiple spatiotemporal features. Since the correlation between different spatiotemporal features and different timescales is often strong, we expected that the intrinsic topological configuration could be faithfully depicted within a reduced-dimensional space. We adopted a semi-supervised framework and leveraged the semi-supervised extension of the Uniform Manifold Approximation and Projection (*90*) algorithm with Hellinger distance. Specifically, given data from an unannotated experiment (i.e., fly), we computed a 2-dimensional embedding by pairing the unannotated experiment with each annotated experiment one at a time. Here, each annotated experiment constitutes a different “view” on the data. When the behavioral repertoire of the annotated and the novel experiment are similar, the provided “view” yields an accurate, easy-to-interpret low-dimensional representation of the exhibited behaviors in the novel experiment. When the behavioral repertoire and/or feature distribution are dissimilar, the resulting embedding may not be informative; however, other paired-embeddings will not be distorted by uninformative or low-quality views.

Following the construction of pairwise embeddings, a nearest-neighbor analysis is deployed on each embedding, to generate a behavior category weight vector for each frame from the unannotated experiment. The entries within the behavioral weight vectors correspond to the similarities with different behavioral categories. Subsequently, an ensemble committee of classifiers is formed for each pairing. Within this committee, each behavioral weight contributes to the ultimate prediction. By aggregating these behavioral weight vectors through summation, followed by the application of L1 normalization, we arrived at final prediction scores that collectively sum up to 1. Subsequently, the final ensemble committee of classifiers aggregates the weights from each pairing for each behavioral category within a frame and performs L1 normalization across the categories. This results in a probabilistic assignment of each unannotated frame within an experiment to one of the five behavioral categories.

### Arousal Threshold

Experiments were performed on 5-7 day old flies, similar to the baseline experiments, with the exception of the chambers. Flies were inserted in a chamber 2.4mm (D) x 4.2mm (H) x 5.6mm (W) at the front and 3.5mm (H) x 5.2mm (W) at the back. Chambers were printed either using a Ultimaker 3 printer or professionally through Protolabs. A coverslip (CS8R, Warner Instruments) coated with Sigmacote was taped in the front and the back of the chamber. A 1064 nm laser diode (L1064H2, Thorlabs) was collimated using an aspheric lens (C240TMD-C, Thorlabs) and expanded using a beam expander (GBE15-C, Thorlabs). A ring light was placed above the fly and aluminum covered plastic was placed to direct the light towards the chamber. To block laser illumination in the video, a notch filter (NF1064-44, Thorlabs) was placed between the chamber and the camera lens.

Experiments started at ZT10 and lasted until ZT0. A custom-written Python script processed the incoming video feed and determined if the fly is moving or not based on a background subtraction algorithm (for details, see https://github.com/mfkeles/flysleepcl). When a fly became immobile for 30 sec, the laser was triggered using a NI-DAQ board, and the power of the laser was linearly increased over 30 sec (fig. S2A). During this perturbation, if a fly started moving, the script ensured that the fly moved at least 90% of the time within a 3-sec window. This ensured that the fly was fully aroused, and the laser would then be turned off. Each perturbation was spaced, so that a perturbed fly would not be perturbed again within a 30 min period.

### Arousal Threshold Data Analysis

For each epoch of perturbation, a latency-to-wake parameter was calculated. To achieve this, the cumulative sum of the calculated motion for each perturbation bout was calculated using the analyzed movement/quiescence data. Then, a breakpoint was calculated where the fly consistently moved using a linearly penalized segmentation algorithm (https://github.com/deepcharles/ruptures). This breakpoint was taken as the arousal point. Flies that spent less than 70% of the time quiescent between ZT13 and ZT23 were discarded. The remaining flies were then scored for the presence or absence of HS behavior during stimulation. This classification was determined by a change in haltere position in the dorsoventral axis that preceded laser-induced movement. Videos corresponding to each arousal event were extracted starting from the closest movement bout defined by 1 sec of continuous movement. Each video was then annotated for HS using three classes: True, False and Inconclusive. Videos were labeled inconclusive when halteres were not visible due to occlusion or body orientation; these data points were excluded from the data analysis. In addition, feeding, grooming, PE, and defecation behaviors that occurred between the last significant movement and laser perturbation were manually annotated using BORIS (*91*). The annotator was blind to determined threshold and other experimental parameters. To calculate the normalized arousal threshold values, the recorded analog voltage signal (copy of the signal sent to the laser driver) was integrated over the calculated point where the animal started moving. For each animal, integrated (Volt-second) values were normalized by subtracting the mean of all the trials per animal and dividing by the standard deviation.

### Optogenetic Activation

Flies with indicated genotypes were loaded into the chambers described above. Experiments were performed between ZT3-ZT9. Flies were acclimated into the chamber for 10 mins. Stimulation protocol consisted of 5 mins of baseline recording, 5 mins of LED stimulation at 1 Hz with a pulse width of 5 ms and finally 10 mins of recording post-stimulation. 1 Hz optogenetic stimulation was chosen because this frequency was within the range observed from recordings of dFB and R5 neurons (*38*, *92*). Camera and LEDs are triggered using an Arduino microcontroller. Two LEDs (590 nm) wired serially were used for stimulation. Resulting videos were manually annotated using BORIS (*91*) and plotted using MATLAB.

### Analysis of FlyVISTA results

We first filtered the behavioral predictions based on the confidence score for the body part of interest that is associated with the targeted behavior. For example, the behavioral score for HS is assigned to zero for frames where DeepLabCut has a threshold less than 0.8 for the location of the haltere facing the camera. We used the same threshold for the position of the proboscis and applied it to PE and feeding behaviors. For grooming, thorax position was used for filtering. We then assigned the behavior with the highest behavioral score to that frame as the designated behavior. The resulting arrays were then further processed to calculate the duration and bout number per behavior as appropriate.

To calculate sleep, we considered grooming, feeding and movement behavioral classes and calculated continuous bouts that excluded those categories. We then further filtered these data by applying our sleep criteria based on arousal threshold experiments: consolidated bouts of quiescence lasting ≥3 min were identified as sleep.

### Representative Traces

Representative traces in Figs. 1, S2G and H, 7J, S1A, S7A-D, were plotted using the data generated after the feature extraction step of FlyVISTA. To calculate power spectral density plots, 2 min windows of oscillatory haltere sections are lowpass filtered at 0.5 Hz with a 5^th^ order Butterworth filter. These traces were then used to calculate periodograms.

### Tracking multiple body parts with CoTracker

To track points not included in our DeepLabCut dataset (i.e., arousal threshold data and antennal base and tip movements), we used CoTracker (*93*), a transformer-based model that allows tracking of any pixel in a video. To achieve this, we first trimmed the portion of the video to be analyzed. The video was then cropped to 512×384 pixels, which is the width and height of the training dataset used in CoTracker (*93*). We then selected the desired number of points on the body of the fly and tracked these points. Traces were low-pass filtered at 15 Hz using a 2^nd^ order Butterworth filter, with the exception of the leg traces shown in Fig. S1C which were bandpass filtered (low = 0.5 Hz, high = 10 Hz). CoTracker was used in the following figures: S1B and C, 2G, S2D-F, S7E and F.

We calculated power spectral density using the following method. For each signal trace in the dataset, we applied zero-padding to the next power of two to increase frequency resolution. We then computed the periodogram using the periodogram function with a sampling frequency of 30 Hz. The resulting power values were normalized for each fly.

### Tracking of freely moving flies during optogenetic activation

To track position of the fly in the arena and generate a segmentation mask we trained a custom state-of-the-art computer vision model YOLOv8 (*94*). We annotated segmentation masks for 570 images derived from the optogenetic activation videos. This dataset was split into training, validation, and test sets. Initially, the dataset was divided into an 80/20 split for training and validation. The validation set was then further split equally to create final validation and test sets, resulting in each of these sets comprising 10% of the total data. We employed a YOLOv8 segmentation model, which was pre-trained and then fine-tuned using our custom dataset. The training process was configured to run for 100 epochs with a batch size of 16 and an image size of 640 pixels. We used an AdamW optimizer with an initial learning rate set at 0.01. The model training was conducted on two GPUs (RTX 3090, NVIDIA). The model achieved a precision of 96.0% and a recall of 84.8% on the validation set. The mean Average Precision (mAP) at 50% Intersection over Union (IoU) was 92.0%. The resulting model was used to track the position of the fly, as well as its segmentation mask. Coordinates for centroid points are first median filtered with a window size of 7. This is followed by calculating the change in position in x and y coordinates. These values are then used to calculate the Euclidian distance between each consecutive points in the time series data. The resulting data is then converted to mm/sec by using the sampling rate of the video and corresponding mm value per pixel. Distance is calculated in a similar fashion where the total distance moved is summed over each minute. To visualize the location of the fly in the arena, data is downsampled so that each data point represented a second. This is then plotted over the corresponding bins as shown in Fig S3G.

### Trikinetics Sleep Monitoring

Sleep analyses using the *Drosophila* Activity Monitoring System (DAM) (Trikinetics, Inc.) were performed as previously described (*73*). In brief, male or female flies were anesthetized with CO_2_ and loaded into wax- and yarn-sealed glass tubes containing standard food. Flies were allowed to recover from CO_2_ for at least 36 hrs, and flies were 5-6 days old during the recording. Experiments were conducted in an incubator at 25°C and ∼60% relative humidity. Sleep was identified using the traditional definition of 5 min consolidated inactivity (*24*) and analyzed using Sleeplab (*95*).

### Sleep Deprivation

Flies were loaded into individual DAM tubes using CO_2_ anesthesia 2 days prior to the video monitoring. Flies were then mechanically sleep-deprived 3 sec every min from ZT12-ZT0 using a vortexer (VWR DVX-2500 Multi-tube vortexer). Flies were then taken out of the individual tubes between ZT23 and ZT0 and placed into standard imaging chambers using a mouth pipette. Video recording started at ZT0 and lasted until ZT6. The ZT0 to ZT2 portion of the video was used for analysis.

### Statistical analysis

For comparisons of 2 groups of normally distributed data, unpaired or paired t-tests were performed. For multiple comparisons of normally distributed data, one-way ANOVAs followed by post-hoc Tukey were performed. For comparisons of 2 groups of non-normally distributed data, Mann-Whitney U-tests were performed. For multiple comparisons of non-normally distributed data, Kruskall-Wallis tests with post-hoc Dunn’s was performed. For comparisons of 2 distributions, Kolgomorov-Smirnov tests were used.

## Acknowledgments

We thank Vivek Jayaraman and the Bloomington Stock Center for flies. We thank Deniz Yavuz for assistance with the laser setup, Deniz Cem Önduygu for illustrating the computational pipeline and Yusufhan Kırçova for assistance with the closed-loop video processing. We thank Anuradha Mehta for assistance with Trikinetics data and William Joiner for Sleeplab.

## Author contributions

Conceptualization: M.F.K, M.N.W. Methodology: M.F.K., A.O.B.S., O.T., S.K.

Investigation: M.F.K., A.O.B.S., C.B., I.P., C.L. Visualization: M.F.K., A.O.B.S.

Supervision: M.F.K., O.T., S.K., M.N.W. Writing (original draft): M.F.K., M.N.W.

Writing (review and editing): M.F.K., A.O.B.S., C.B., I.P., C.L., O.T., S.K., M.N.W.

## Data and materials availability

https://mfkeles.github.io/FlyVISTA/

## Supplemental Figure Legends

**Supplementary Figure 1:**
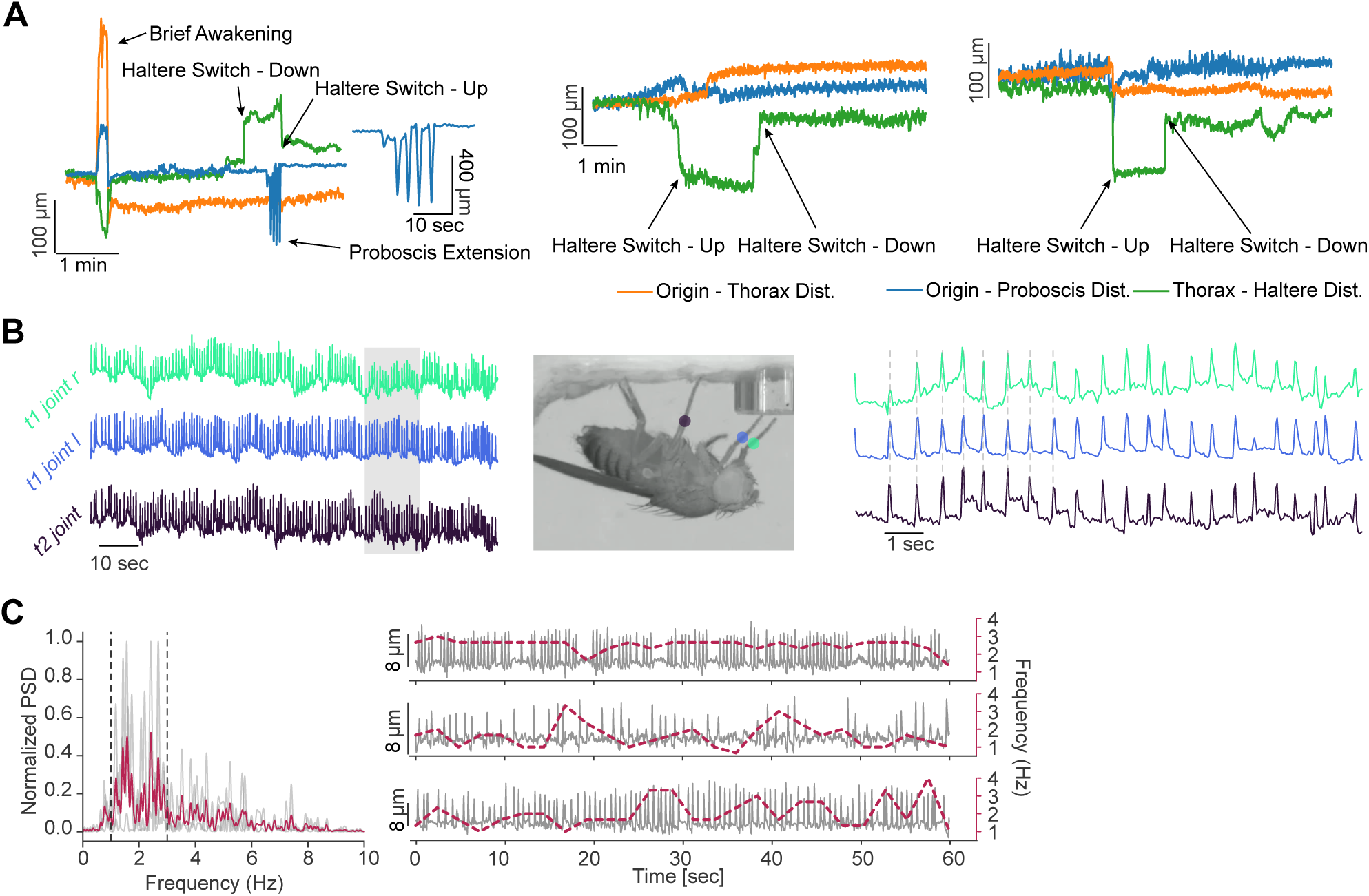
Varying spatiotemporal features of microbehaviors observed during quiescence. (**A**) Additional examples of quiescence-associated behaviors, showing spatiotemporal structure of HS and PE. Orange, magenta, and green lines indicate the distance between origin (fixed point at 0, 0) and thorax, origin and proboscis tip, and thorax to haltere, respectively. “Haltere Switch – Down” refers to HS towards the ventral direction, while “Haltere Switch – Up” refers to HS towards the dorsal direction. Expanded trace of PE is shown in the left plot. **(B)** Periodic movement of the legs is fast and synchronized across legs. Time series of tracked points are shown (left) along with a representative image (middle, colors on the representative image correspond to the colors of the time series data). Right panel, expanded time series data showing coordinated fast movement of the legs, with the peaks of all three traces aligning across the dashed gray lines. **(C)** Normalized power spectral density plots show the frequency of the rapid leg movements are stereotyped across flies (n=3). Mean is shown in red. Dashed lines indicate 1 to 3 Hz borders. Right panel shows 3 sec window max Short-time Fourier Transform (dashed red line) overlaid on the distance between a fixed point and a tracked leg from an individual fly (gray line).

**Supplementary Figure 2:**
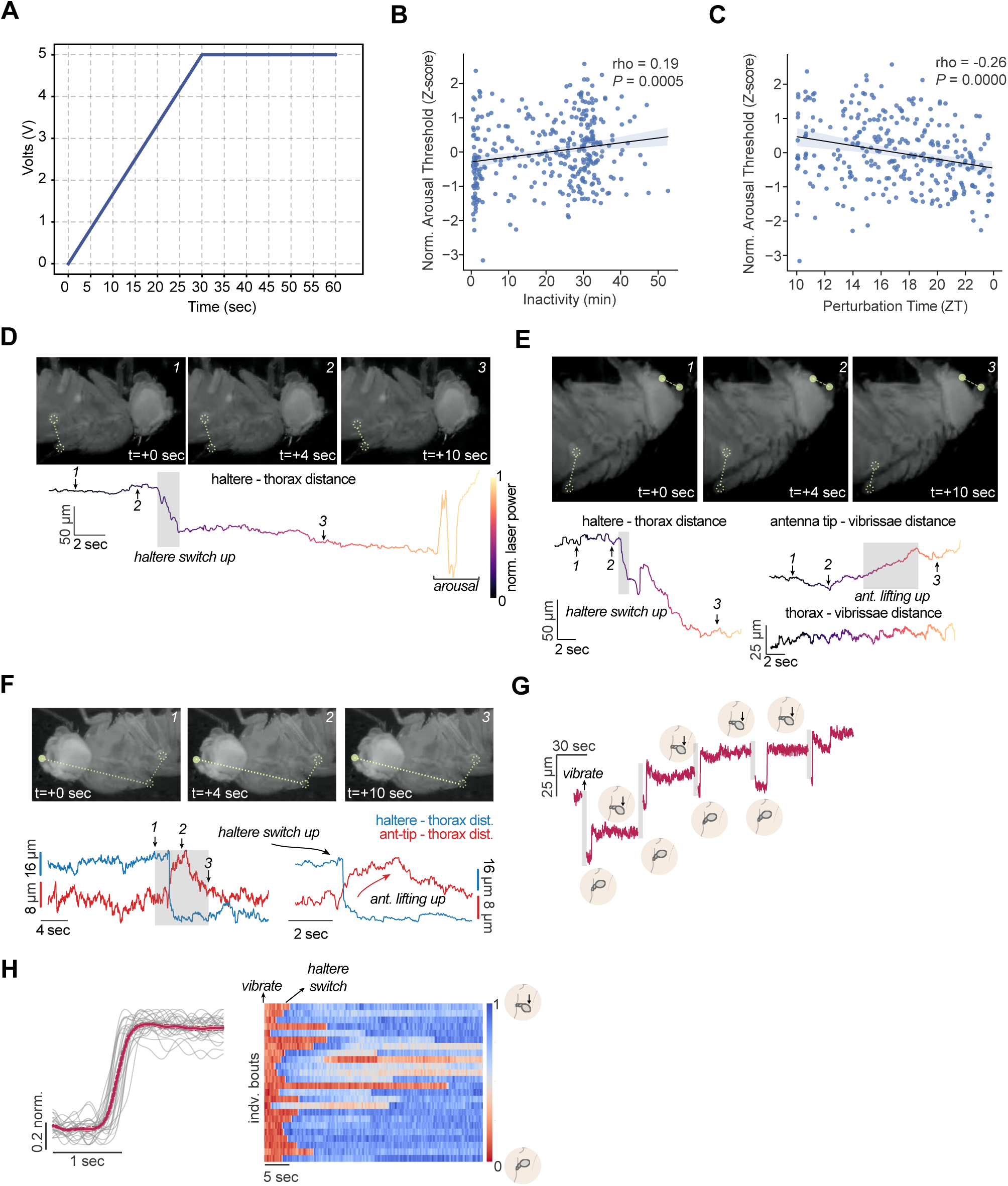
Additional information related to arousal threshold experiments. (**A**) Perturbation protocol for assessing arousal threshold. Plot illustrating voltage/time relationship, where the laser is turned on and its power is linearly increased over 30 sec. (**B**) Correlation between normalized arousal thresholds (Z-score) for individual perturbation bouts vs inactivity bout duration for female flies (n=16). Linear regression shows positive correlation between arousal threshold and increased quiescence bout duration. (**C**) Correlation between normalized arousal thresholds (Z-score) for individual perturbation bouts vs ZT time for female flies, where the bouts were derived from events where the animal exhibited movement in the 30-60 sec window prior to “laser-on.” Linear regression shows negative correlation between arousal threshold and ZT time. **(D** and **E**) Representative images (above) accompanied by time series for the tracked points (below, haltere and scutellum or antenna and vibrissae, dashed circles) show changes in microbehaviors (HS up and/or antennal movement upward) that can occur prior to arousal. Shaded boxes denote HS up and antennal lifting up. Varying color on the time series data indicates the change in the laser power. **(F)** Representative images (above) accompanied by time series data (below) for the tracked points (haltere and scutellum, dashed circles; antenna tip, solid circle). Simultaneous occurrence of antennal movement (red) and HS (blue) during a spontaneous sleep bout. An expanded time series (bottom right) is shown corresponding to the shaded box in the bottom left time series. **(G)** Representative HS events following individual perturbation bouts in an individual fly. Distance of the haltere to a fixed point plotted in red. Shaded areas indicate where the vibration motor is turned on. Cartoon illustrations indicate the haltere position. **(H)** Time aligned, normalized HS events after each perturbation bout (left). Solid line and dashed lines indicate mean and SEM, respectively. Heatmap (right) of the individual bouts (n=24) and the change in the haltere position (red to blue) is shown in each bout.

**Supplementary Figure 3:**
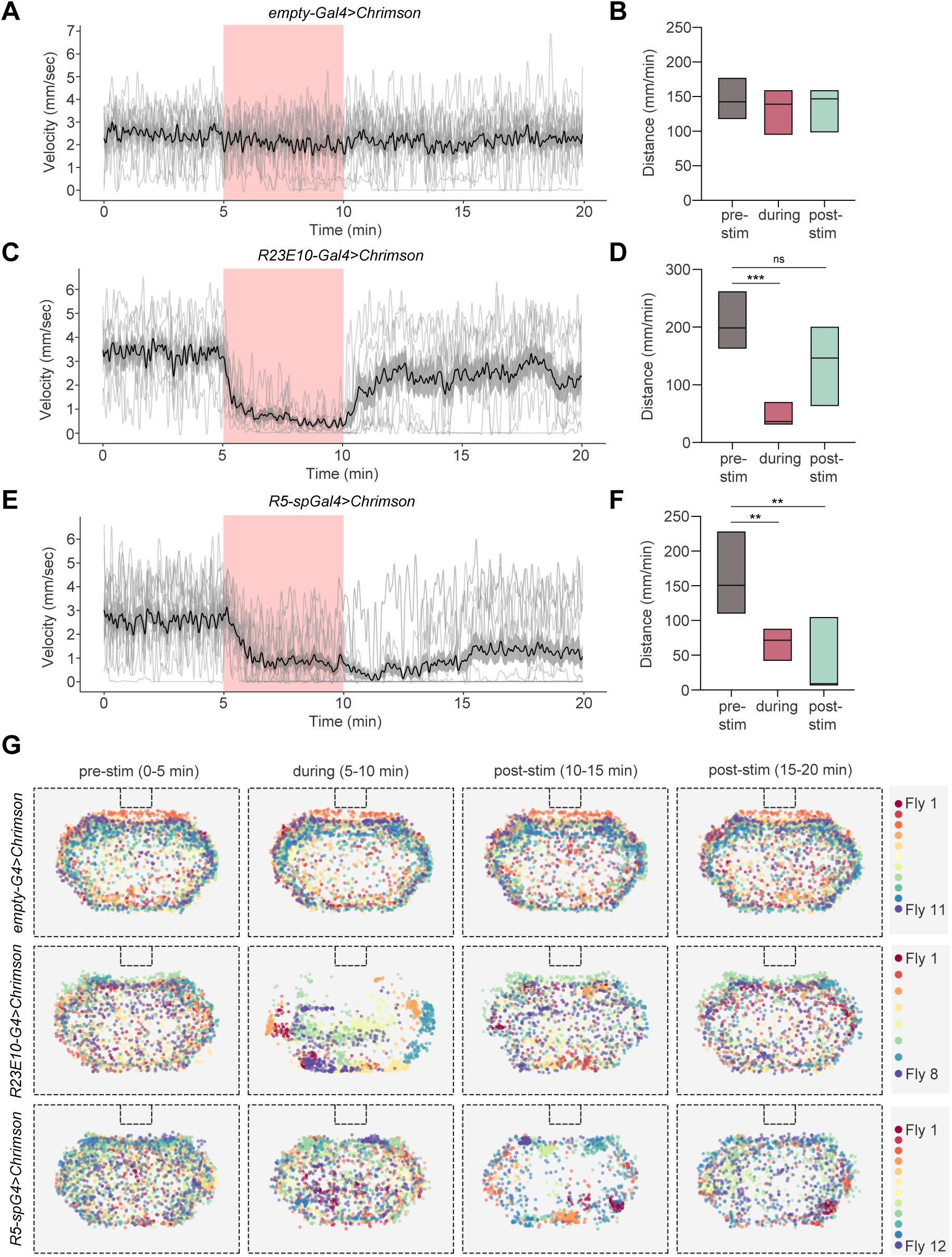
Effects of optogenetic activation of sleep-related circuits. **(A, C, and E)** Velocity of individual (gray) and mean (black) flies for the 3 genotypes shown over a 20 min period with optogenetic activation indicated by the red box. **(B, D,** and **F)** Simplified box plots showing the distance covered (mm/min) by each group before (“pre-stim”), during (“during”), and after (“post-stim”) Chrimson activation. Kruskall-Wallis test, with post-hoc Dunn’s test. ns, not significant; **, *P*<0.01; and ***, *P*<0.001. **(G)** Position of the centroid in the arena for each tracked fly across different genotypes, before, during, and after optogenetic stimulation. Outer dashed lines indicate the border of the imaging view and the top inner dashed rectangle indicate the location of the food port. Each fly has a distinct color, and centroid position is plotted in 1 sec intervals. Data in this figure are from the same flies as in Fig. 3.

**Supplementary Figure 4:**
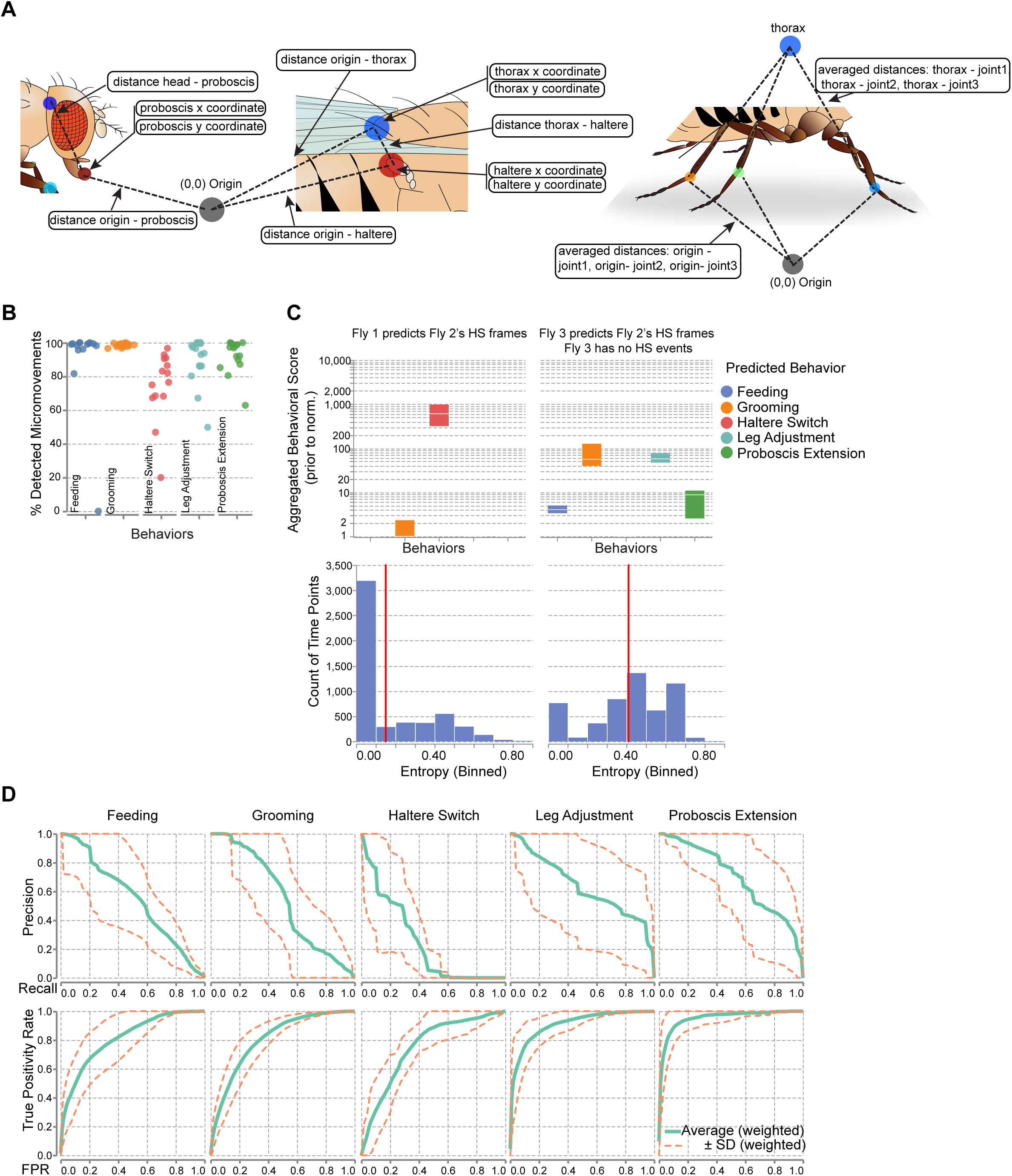
Performance metrics and details for the FlyVISTA computational pipeline. (**A**) Schematics illustrating the body parts or X-Y location (origin) used in the 13 features for the computational pipeline. (**B**) Evaluation of the micro-activity detection stage of the pipeline. Each data point stands for a leave-one-out experiment, and the *y*-axis represents the percentage of time points correctly detected as micro-activity for each behavior category. Low percentage values are undesired, as only the time points detected as micro-activity are analyzed in the latter stages of the pipeline and directly affect the downstream analysis. Extremely low percentages often indicate behaviors that are rarely exhibited, rather than poor detection performance. (**C**) Simplified box-plots (75^th^ percentile, median, 25^th^ percentile) of behavioral scores before normalization (above) and histograms of corresponding entropy values (below) are computed using one unannotated (Fly 2) and two annotated (Fly 1 and Fly 3) experiments with varying behavioral repertoires. Behavioral scores and entropy values are computed for the HS behavior. Fly 1 exhibited HS behavior, whereas Fly 3 did not. As a result, behavioral scores generated by comparing to Fly 3 fail to indicate a confident behavioral category, and hence, tend to have higher entropy (right lower panel). In contrast, confident predictions and low entropy of behavioral score distributions for HS behavior are observed with Fly 1’s annotations (left sub-figure). These results demonstrate the ability of the computational pipeline to detect and discover unseen unannotated behavioral categories using behavioral scores. (**D**) Performance summary of behavior mapping demonstrated using receiver operating characteristic curve and precision-recall curve. The weighted averages (solid lines) and standard deviations (dashed lines) of ROC and precision-recall curves are computed from all sixteen leave-one-out by interpolation.

**Supplementary Figure 5:**
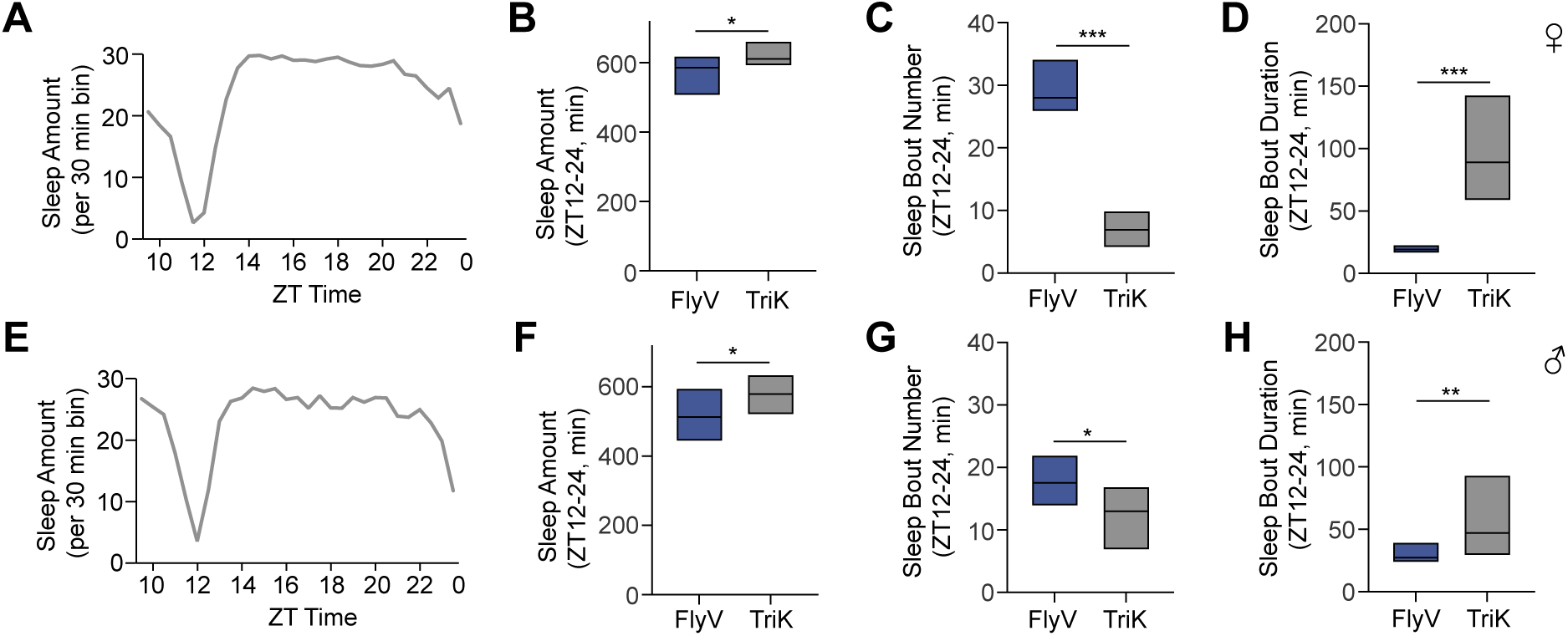
Comparison of Trikinetics and FlyVISTA data. (**A** and **E**) Sleep profiles from ZT10 to ZT0 measured using the Trikinetics system for female (n=31) (**A**) and male (n=34) (**E**) flies. (**B**-**D**) Nighttime sleep amount (**B**), sleep bout number (**C**), and sleep bout duration (**D**) for female flies using FlyVISTA (FlyV) vs Trikinetics (TriK) DAM monitors. (**F**-**H**) Nighttime sleep amount (**F**), sleep bout number (**G**), and sleep bout duration (**H**) for male flies using FlyV vs TriK DAM monitors. FlyV data are from the flies shown in Figs. 5A-5C. Simplified box plots are shown in **B**-**D** and **F**-**H**, where top and bottom of the box denote 75^th^ and 25^th^ percentiles, and middle line indicates median. Mann Whitney U-test; *, *P* < 0.05; **, *P* < 0.01; ***, *P* < 0.001.

**Supplementary Figure 6:**
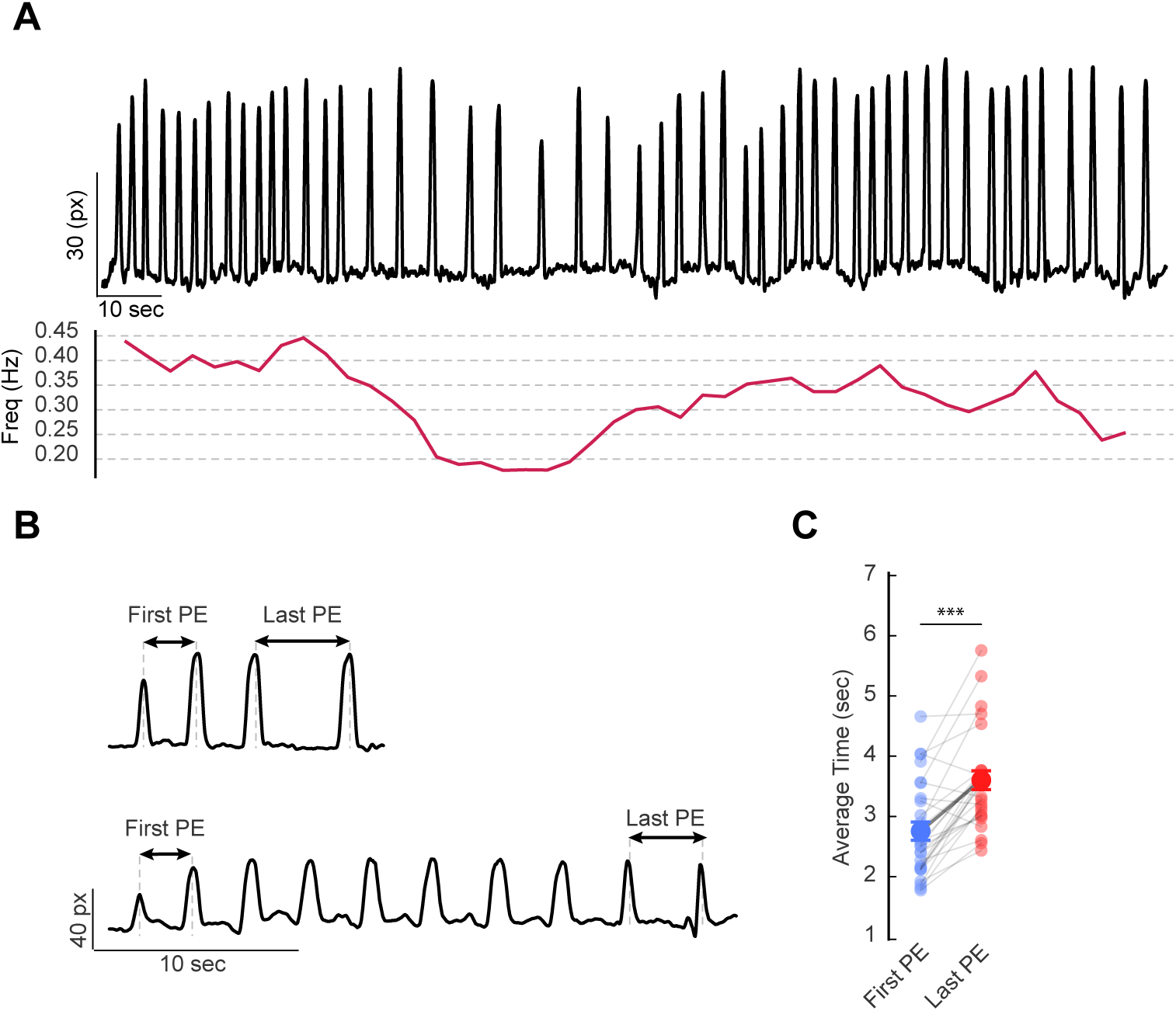
Spatiotemporal structure of PE microbehavior. (**A**) Example trace (above, origin-proboscis distance) and PE frequency (below, in 3 sec bins) showing variability of the inter-PE interval within a single long bout from a male fly. (**B**) 2 example traces (origin-proboscis distance) of individual PE bouts showing a longer inter-PE interval for the last 2 PEs compared to the first 2 PEs. (**C**) Quantification of the first and last inter-PE intervals under baseline conditions for the animals shown in Fig. 6A, with male and females pooled. n=28, paired t-test; ***, *P* < 0.001.

**Supplementary Figure 7:**
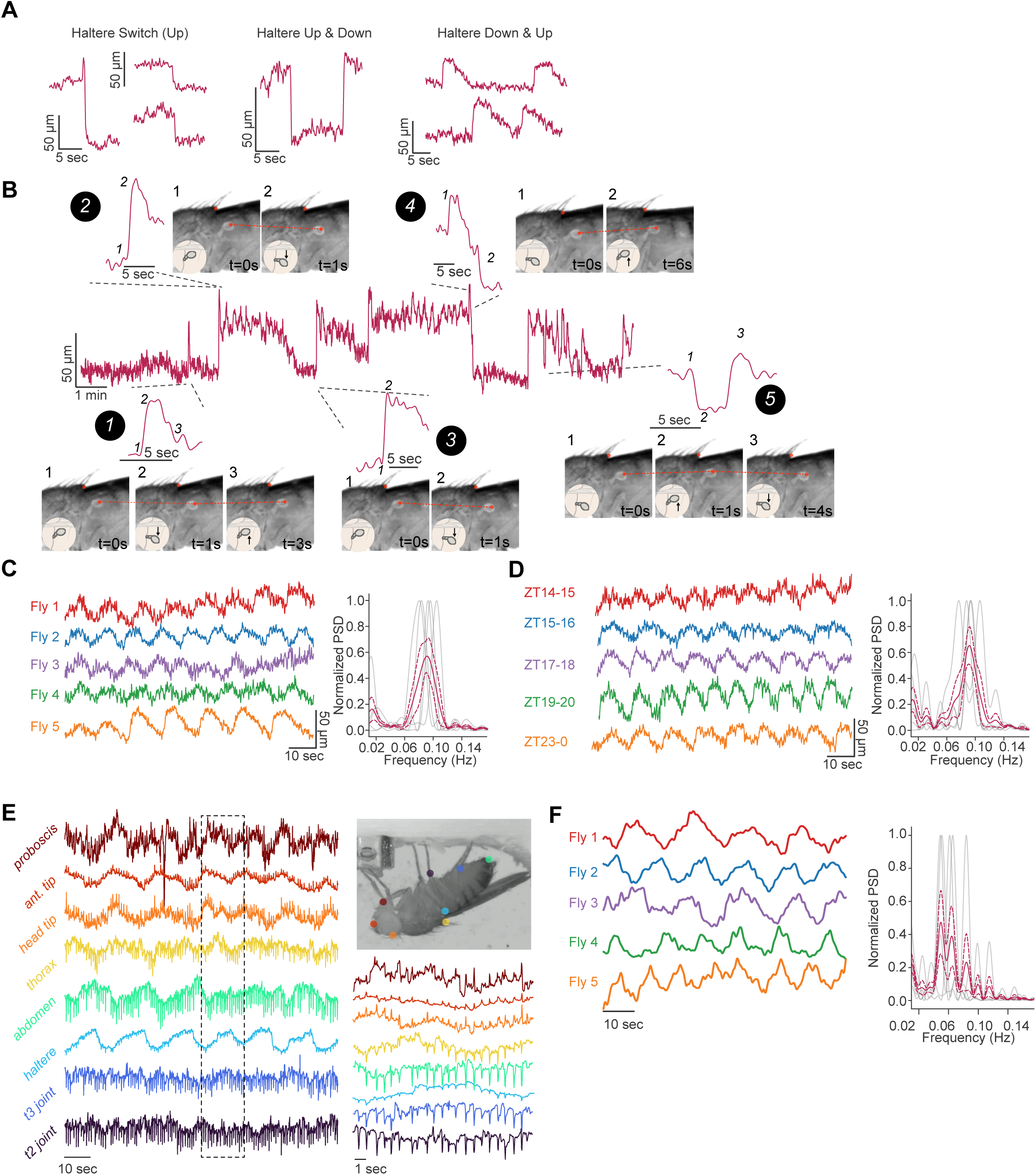
HS microbehaviors exhibit diverse phenotypes. (**A**) Example time traces of a variety of haltere switch behaviors with distinct spatiotemporal organization. Haltere Switch (Up) behavior is marked by a sudden change in the distance between halteres and the most dorsoposterior part of the scutellum. Haltere up and down behavior reflects a change in the haltere position towards the dorsal side followed by a change towards the ventral side in a rapid fashion. Haltere Down and Up behavior occurs when halteres move down towards the abdomen but return to their initial position gradually. (**B**) Example time trace of active haltere behavior illustrated by the dynamic changes in the distance between halteres and scutellum over ∼18 mins. Individual HS events are shown above or below the main time trace with the accompanying changes in the haltere position indicated by expanded time traces (left) and representative images (right). Behaviors are highlighted in the following order (numbers in solid circles): 1) Haltere down & up, 2) Haltere down, 3) Haltere down, 4) Haltere up, 5) Haltere up & down. Dashed lines indicate the location of the expanded traces. Cartoons show the changes in the haltere position with consecutive time points. **(C** and **D)** Oscillatory movement of the halteres across different individuals (**C**) and across different ZT times (**D**). 1 min examples of the haltere position from 5 flies or 5 ZT times (indicated by different colors) are shown. Traces (left) indicate the distance of haltere position to a fixed point. Normalized power spectral density (PSD, right) is shown. (**E)** Oscillations across tracked points along the body. 8 tracked points suggest oscillations across the body are synchronized. Tracked points on the fly body are shown in top right. Expanded traces of the dashed box for each tracked point are shown (bottom right). **(F)** Traces indicate the distance of abdomen to a fixed point for 5 flies. Normalized PSD (right) is shown. For PSD plots, individual data (gray), mean (red line), and SEM (dashed lines) are shown.

**Supplementary Video 1: Rhythmic proboscis extension behavior.** Video of wild-type fly exhibiting proboscis extension behavior during quiescence. Time series trace on the right indicates the change in distance between the tracked point and a fixed point.

**Supplementary Video 2: Haltere switch behavior.** Video of wild-type fly exhibiting haltere switch behavior. Time series traces on the right indicate changes in position of tracked points (red, haltere; blue, scutellum). Arrow indicates onset of HS “down” behavior.

**Supplementary Video 3: Optogenetic activation using *R23E10-Gal4* to target dFB neurons.** Four representative videos span 10 sec before and 45 sec during “LED ON” and show frequent grooming behavior.

**Supplementary Video 4: Optogenetic activation using *R23E10-Gal4* to target dFB neurons.** Four representative videos span 20 sec during “LED ON” and show frequent micromovements.

**Supplementary Video 5: Optogenetic activation using *R58H05-AD, R46C03-DBD* to target R5 neurons.** Four representative videos span 10 sec before and 1 min during “LED ON” and reveal proboscis extension behavior.

**Supplementary Video 6: Optogenetic activation using *R58H05-AD, R46C03-DBD* to target R5 neurons.** Four representative videos span 10 sec before and 2 min after “LED OFF” and highlight the transition into a quiescent state. For videos S3-S6, behavior is noted in the top right corner. For videos S3, S5, and S6, “LED ON” and “LED OFF” is noted in the top left corner. Time stamps are shown in the top middle.

## References

1. S. W. Flavell, N. Gogolla, M. Lovett-Barron, M. Zelikowsky, The emergence and influence of internal states. Neuron 110, 2545–2570 (2022).

2. D. J. Anderson, P. Perona, Toward a science of computational ethology. Neuron 84, 18–31 (2014).

3. S. E. R. Egnor, K. Branson, Computational Analysis of Behavior. Annu. Rev. Neurosci. 39, 217–236 (2016).

4. S. R. Datta, D. J. Anderson, K. Branson, P. Perona, A. Leifer, Computational Neuroethology: A Call to Action. Neuron 104, 11–24 (2019).

5. T. D. Pereira, J. W. Shaevitz, M. Murthy, Quantifying behavior to understand the brain. Nat. Neurosci., 1–13 (2020).

6. N. Tinbergen, Comparative Studies of the Behaviour of Gulls (Laridae): A Progress Report. Behaviour 15, 1–70 (1959).

7. C. Darwin, P. Ekman, The Expression of the Emotions in Man and Animals, Definitive Edition (Oxford University Press, ed. 3, 1998; https://www.amazon.com/Expression-Emotions-Man-Animals-Definitive/dp/0195112717).

8. A. Mathis, P. Mamidanna, K. M. Cury, T. Abe, V. N. Murthy, M. W. Mathis, M. Bethge, DeepLabCut: markerless pose estimation of user-defined body parts with deep learning. Nat. Neurosci. 21, 1281–1289 (2018).

9. R. Allada, J. M. Siegel, Unearthing the phylogenetic roots of sleep. Curr. Biol. 18, R670–R679 (2008).

10. G. Oikonomou, D. A. Prober, Attacking sleep from a new angle: contributions from zebrafish. Curr. Opin. Neurobiol. 44, 80–88 (2017).

11. R. C. Anafi, M. S. Kayser, D. M. Raizen, Exploring phylogeny to find the function of sleep. Nat. Rev. Neurosci. 20, 109–116 (2019).

12. T. L. Iglesias, J. G. Boal, M. G. Frank, J. Zeil, R. T. Hanlon, Cyclic nature of the REM sleep-like state in the cuttlefish *Sepia officinalis*. J. Exp. Biol. 222 (2019).

13. R. D. Nath, C. N. Bedbrook, M. J. Abrams, T. Basinger, J. S. Bois, D. A. Prober, P. W. Sternberg, V. Gradinaru, L. Goentoro, The Jellyfish *Cassiopea* Exhibits a Sleep-like State. Curr. Biol. 27, 2984–2990.e3 (2017).

14. I. Tobler I., M. Neuner-Jehle, 24-h variation of vigilance in the cockroach Blaberus giganteus. J. Sleep Res. 1, 231–239 (1992).

15. D. M. Raizen, J. E. Zimmerman, M. H. Maycock, U. D. Ta, Y.-J. You, M. V. Sundaram, A. I. Pack, Lethargus is a *Caenorhabditis elegans* sleep-like state. Nature 451, 569–572 (2008).

16. S. L. de S. Medeiros, M. M. M. de Paiva, P. H. Lopes, W. Blanco, F. D. de Lima, J. B. C. de Oliveira, I. G. Medeiros, E. B. Sequerra, S. de Souza, T. S. Leite, S. Ribeiro, Cyclic alternation of quiet and active sleep states in the octopus. iScience 24, 102223 (2021).

17. T. Yokogawa, W. Marin, J. Faraco, G. Pézeron, L. Appelbaum, J. Zhang, F. Rosa, P. Mourrain, E. Mignot, Characterization of sleep in zebrafish and insomnia in hypocretin receptor mutants. PLoS Biol. 5, e277 (2007).

18. A. Rechtschaffen, M. A. Gilliland, B. M. Bergmann, J. B. Winter, Physiological correlates of prolonged sleep deprivation in rats. Science 221, 182–184 (1983).

19. G. Tononi, C. Cirelli, Sleep and the price of plasticity: from synaptic and cellular homeostasis to memory consolidation and integration. Neuron 81, 12–34 (2014).

20. V. M. Hill, R. M. O’Connor, M. Shirasu-Hiza, Tired and stressed: Examining the need for sleep. Eur. J. Neurosci. 51, 494–508 (2020).

21. L. Imeri, M. R. Opp, How (and why) the immune system makes us sleep. Nat. Rev. Neurosci. 10, 199–210 (2009).

22. E. Mignot, Why we sleep: the temporal organization of recovery. PLoS Biol. 6, e106 (2008).

23. L. Xie, H. Kang, Q. Xu, M. J. Chen, Y. Liao, M. Thiyagarajan, J. O’Donnell, D. J. Christensen, C. Nicholson, J. J. Iliff, T. Takano, R. Deane, M. Nedergaard, Sleep drives metabolite clearance from the adult brain. Science 342, 373–377 (2013).

24. P. J. Shaw, C. Cirelli, R. J. Greenspan, G. Tononi, Correlates of sleep and waking in *Drosophila melanogaster*. Science 287, 1834–1837 (2000).

25. J. C. Hendricks, S. M. Finn, K. A. Panckeri, J. Chavkin, J. A. Williams, A. Sehgal, A. I. Pack, Rest in *Drosophila* is a sleep-like state. Neuron 25, 129–138 (2000).

26. J. E. Zimmerman, N. Naidoo, D. M. Raizen, A. I. Pack, Conservation of sleep: insights from non-mammalian model systems. Trends Neurosci. 31, 371–376 (2008).

27. J. C. Chiu, K. H. Low, D. H. Pike, E. Yildirim, I. Edery, Assaying locomotor activity to study circadian rhythms and sleep parameters in *Drosophila*. J. Vis. Exp., doi: 10.3791/2157 (2010).

28. M. E. Driscoll, C. Hyland, D. Sitaraman, Measurement of Sleep and Arousal in *Drosophila*. Bio Protoc 9 (2019).

29. G. F. Gilestro, Video tracking and analysis of sleep in *Drosophila melanogaster*. Nat. Protoc. 7, 995–1007 (2012).

30. N. C. Donelson, E. Z. Kim, J. B. Slawson, C. G. Vecsey, R. Huber, L. C. Griffith, High-resolution positional tracking for long-term analysis of *Drosophila* sleep and locomotion using the “tracker” program. PLoS One 7, e37250 (2012).

31. E. Aserinsky, N. Kleitman, Regularly occurring periods of eye motility, and concomitant phenomena, during sleep. Science 118, 273–274 (1953).

32. J. A. Hobson, T. Spagna, R. Malenka, Ethology of sleep studied with time-lapse photography: postural immobility and sleep-cycle phase in humans. Science 201, 1251–1253 (1978).

33. W. Kaiser, Busy bees need rest, too. Journal of Comparative Physiology A 163, 565–584 (1988).

34. S. Sauer, M. Kinkelin, E. Herrmann, W. Kaiser, The dynamics of sleep-like behaviour in honey bees. J. Comp. Physiol. A Neuroethol. Sens. Neural Behav. Physiol. 189, 599–607 (2003).

35. B. van Alphen, E. R. Semenza, M. Yap, B. van Swinderen, R. Allada, A deep sleep stage in *Drosophila* with a functional role in waste clearance. Science Advances 7, eabc2999 (2021).

36. J. M. Donlea, M. S. Thimgan, Y. Suzuki, L. Gottschalk, P. J. Shaw, Inducing sleep by remote control facilitates memory consolidation in *Drosophila*. Science 332, 1571–1576 (2011).

37. M. C. W. Ho, M. Tabuchi, X. Xie, M. P. Brown, S. Luu, S. Wang, A. L. Kolodkin, S. Liu, M. N. Wu, Sleep need-dependent changes in functional connectivity facilitate transmission of homeostatic sleep drive. Curr. Biol. 32, 4957–4966.e5 (2022).

38. S. Liu, Q. Liu, M. Tabuchi, M. N. Wu, Sleep Drive Is Encoded by Neural Plastic Changes in a Dedicated Circuit. Cell 165, 1347–1360 (2016).

39. G. Fraenkel, J. W. S. Pringle, Biological Sciences: Halteres of Flies as Gyroscopic Organs of Equilibrium. Nature 141, 919–920 (1938).

40. K. A. Daltorio, J. L. Fox, Haltere removal alters responses to gravity in standing flies. J. Exp. Biol. 221 (2018).

41. H. Piéron, Le problème physiologique du sommeil (E. Grevin, 1913; https://play.google.com/store/books/details?id=kbIxOE7bQ1IC).

42. S. S. Campbell, I. Tobler, Animal sleep: a review of sleep duration across phylogeny. Neurosci. Biobehav. Rev. 8, 269–300 (1984).

43. Q. Geissmann, L. Garcia Rodriguez, E. J. Beckwith, A. S. French, A. R. Jamasb, G. F. Gilestro, Ethoscopes: An open platform for high-throughput ethomics. PLoS Biol. 15, e2003026 (2017).

44. R. Allada, B. Y. Chung, Circadian organization of behavior and physiology in *Drosophila*. Annu. Rev. Physiol. 72, 605–624 (2010).

45. C. Helfrich-Förster, Differential control of morning and evening components in the activity rhythm of *Drosophila melanogaster*--sex-specific differences suggest a different quality of activity. J. Biol. Rhythms 15, 135–154 (2000).

46. C.-L. Chen, F. Aymanns, R. Minegishi, V. D. V. Matsuda, N. Talabot, S. Günel, B. J. Dickson, P. Ramdya, Ascending neurons convey behavioral state to integrative sensory and action selection brain regions. Nat. Neurosci. 26, 682–695 (2023).

47. F.-O. Lehmann, N. Heymann, Unconventional mechanisms control cyclic respiratory gas release in flying *Drosophila*. J. Exp. Biol. 208, 3645–3654 (2005).

48. G. Pass, Accessory pulsatile organs: evolutionary innovations in insects. Annu. Rev. Entomol. 45, 495–518 (2000).

49. C. R. Sharkey, J. Blanco, M. M. Leibowitz, D. Pinto-Benito, T. J. Wardill, The spectral sensitivity of *Drosophila* photoreceptors. Sci. Rep. 10, 18242 (2020).

50. O. T. Shafer, A. C. Keene, The Regulation of *Drosophila* Sleep. Curr. Biol. 31, R38–R49 (2021).

51. B. van Alphen, M. H. W. Yap, L. Kirszenblat, B. Kottler, B. van Swinderen, A dynamic deep sleep stage in *Drosophila*. J. Neurosci. 33, 6917–6927 (2013).

52. L. A. L. Tainton-Heap, L. C. Kirszenblat, E. T. Notaras, M. J. Grabowska, R. Jeans, K. Feng, P. J. Shaw, B. van Swinderen, A Paradoxical Kind of Sleep in *Drosophila melanogaster*. Curr. Biol., doi: 10.1016/j.cub.2020.10.081 (2020).

53. R. Faville, B. Kottler, G. J. Goodhill, P. J. Shaw, B. van Swinderen, How deeply does your mutant sleep? Probing arousal to better understand sleep defects in *Drosophila*. Sci. Rep. 5, 8454 (2015).

54. M. H. W. Yap, M. J. Grabowska, C. Rohrscheib, R. Jeans, M. Troup, A. C. Paulk, B. van Alphen, P. J. Shaw, B. van Swinderen, Oscillatory brain activity in spontaneous and induced sleep stages in flies. Nat. Commun. 8, 1815 (2017).

55. J. M. Donlea, D. Pimentel, C. B. Talbot, A. Kempf, J. J. Omoto, V. Hartenstein, G. Miesenböck, Recurrent Circuitry for Balancing Sleep Need and Sleep. Neuron 97, 378–389.e4 (2018).

56. B. K. Hulse, H. Haberkern, R. Franconville, D. Turner-Evans, S.-Y. Takemura, T. Wolff, M. Noorman, M. Dreher, C. Dan, R. Parekh, A. M. Hermundstad, G. M. Rubin, V. Jayaraman, A connectome of the *Drosophila* central complex reveals network motifs suitable for flexible navigation and context-dependent action selection. Elife 10 (2021).

57. J. D. Jones, B. L. Holder, K. R. Eiken, A. Vogt, A. I. Velarde, A. J. Elder, J. A. McEllin, S. Dissel, Regulation of sleep by cholinergic neurons located outside the central brain in *Drosophila*. PLoS Biol. 21, e3002012 (2023).

58. J. De, M. Wu, V. Lambatan, Y. Hua, W. J. Joiner, Re-examining the role of the dorsal fan-shaped body in promoting sleep in *Drosophila*. Curr. Biol. 33, 3660–3668.e4 (2023).

59. A. S. French, Q. Geissmann, E. J. Beckwith, G. F. Gilestro, Sensory processing during sleep in *Drosophila melanogaster*. Nature, 1–4 (2021).

60. J. Redmon, S. Divvala, R. Girshick, A. Farhadi, You Only Look Once: Unified, Real-Time Object Detection, arXiv [cs.CV] (2015). http://arxiv.org/abs/1506.02640.

61. I. D. Blum, M. F. Keleş, E.-S. Baz, E. Han, K. Park, S. Luu, H. Issa, M. Brown, M. C. W. Ho, M. Tabuchi, S. Liu, M. N. Wu, Astroglial Calcium Signaling Encodes Sleep Need in *Drosophila*. Curr. Biol. 31, 150–162.e7 (2021).

62. L. K. Satterfield, J. De, M. Wu, T. Qiu, W. J. Joiner, Inputs to the sleep homeostat originate outside the brain. J. Neurosci., doi: 10.1523/JNEUROSCI.2113-21.2022 (2022).

63. T. D. Pereira, N. Tabris, A. Matsliah, D. M. Turner, J. Li, S. Ravindranath, E. S. Papadoyannis, E. Normand, D. S. Deutsch, Z. Y. Wang, G. C. McKenzie-Smith, C. C. Mitelut, M. D. Castro, J. D’Uva, M. Kislin, D. H. Sanes, S. D. Kocher, S. S.-H. Wang, A. L. Falkner, J. W. Shaevitz, M. Murthy, SLEAP: A deep learning system for multi-animal pose tracking. Nat. Methods 19, 486–495 (2022).

64. S. R. Jagannathan, T. Jeans, M. N. Van De Poll, B. van Swinderen, Multivariate classification of multichannel long-term electrophysiology data identifies different sleep stages in fruit flies. Sci Adv 10, eadj4399 (2024).

65. S. Riva, J. I. Ispizua, M. T. Breide, S. Polcowñuk, J. R. Lobera, M. F. Ceriani, S. Risau-Gusman, D. L. Franco, Mating disrupts morning anticipation in *Drosophila melanogaster* females. PLoS Genet. 18, e1010258 (2022).

66. A. A. Borbély, A two process model of sleep regulation. Hum. Neurobiol. 1, 195– 204 (1982).

67. M. Tögel, G. Pass, A. Paululat, The *Drosophila* wing hearts originate from pericardial cells and are essential for wing maturation. Dev. Biol. 318, 29–37 (2008).

68. L. K. Scheffer, C. S. Xu, M. Januszewski, Z. Lu, S.-Y. Takemura, K. J. Hayworth, G. B. Huang, K. Shinomiya, J. Maitlin-Shepard, S. Berg, J. Clements, P. M. Hubbard, W. T. Katz, L. Umayam, T. Zhao, D. Ackerman, T. Blakely, J. Bogovic, T. Dolafi, D. Kainmueller, T. Kawase, K. A. Khairy, L. Leavitt, P. H. Li, L. Lindsey, N. Neubarth, D. J. Olbris, H. Otsuna, E. T. Trautman, M. Ito, A. S. Bates, J. Goldammer, T. Wolff, R. Svirskas, P. Schlegel, E. Neace, C. J. Knecht, C. X. Alvarado, D. A. Bailey, S. Ballinger, J. A. Borycz, B. S. Canino, N. Cheatham, M. Cook, M. Dreher, O. Duclos, B. Eubanks, K. Fairbanks, S. Finley, N. Forknall, A. Francis, G. P. Hopkins, E. M. Joyce, S. Kim, N. A. Kirk, J. Kovalyak, S. A. Lauchie, A. Lohff, C. Maldonado, E. A. Manley, S. McLin, C. Mooney, M. Ndama, O. Ogundeyi, N. Okeoma, C. Ordish, N. Padilla, C. M. Patrick, T. Paterson, E. E. Phillips, E. M. Phillips, N. Rampally, C. Ribeiro, M. K. Robertson, J. T. Rymer, S. M. Ryan, M. Sammons, A. K. Scott, A. L. Scott, A. Shinomiya, C. Smith, K. Smith, N. L. Smith, M. A. Sobeski, A. Suleiman, J. Swift, S. Takemura, I. Talebi, D. Tarnogorska, E. Tenshaw, T. Tokhi, J. J. Walsh, T. Yang, J. A. Horne, F. Li, R. Parekh, P. K. Rivlin, V. Jayaraman, M. Costa, G. S. Jefferis, K. Ito, S. Saalfeld, R. George, I. A. Meinertzhagen, G. M. Rubin, H. F. Hess, V. Jain, S. M. Plaza, A connectome and analysis of the adult *Drosophila* central brain. Elife 9 (2020).

69. C. Cirelli, D. Bushey, S. Hill, R. Huber, R. Kreber, B. Ganetzky, G. Tononi, Reduced sleep in *Drosophila Shaker* mutants. Nature 434, 1087–1092 (2005).

70. K. Koh, W. J. Joiner, M. N. Wu, Z. Yue, C. J. Smith, A. Sehgal, Identification of SLEEPLESS, a sleep-promoting factor. Science 321, 372–376 (2008).

71. N. Stavropoulos, M. W. Young, insomniac and Cullin-3 regulate sleep and wakefulness in *Drosophila*. Neuron 72, 964–976 (2011).

72. D. Rogulja, M. W. Young, Control of sleep by cyclin A and its regulator. Science 335, 1617–1621 (2012).

73. S. Liu, A. Lamaze, Q. Liu, M. Tabuchi, Y. Yang, M. Fowler, R. Bharadwaj, J. Zhang, J. Bedont, S. Blackshaw, T. E. Lloyd, C. Montell, A. Sehgal, K. Koh, M. N. Wu, WIDE AWAKE mediates the circadian timing of sleep onset. Neuron 82, 151– 166 (2014).

74. H. Toda, J. A. Williams, M. Gulledge, A. Sehgal, A sleep-inducing gene, nemuri, links sleep and immune function in *Drosophila*. Science 363, 509–515 (2019).

75. T. W. Dunn, J. D. Marshall, K. S. Severson, D. E. Aldarondo, D. G. C. Hildebrand, S. N. Chettih, W. L. Wang, A. J. Gellis, D. E. Carlson, D. Aronov, W. A. Freiwald, F. Wang, B. P. Ölveczky, Geometric deep learning enables 3D kinematic profiling across species and environments. Nat. Methods 18, 564–573 (2021).

76. S. Günel, H. Rhodin, D. Morales, J. Campagnolo, P. Ramdya, P. Fua, DeepFly3D, a deep learning-based approach for 3D limb and appendage tracking in tethered, adult *Drosophila*. Elife 8 (2019).

77. M. Kabra, A. A. Robie, M. Rivera-Alba, S. Branson, K. Branson, JAABA: interactive machine learning for automatic annotation of animal behavior. Nat. Methods 10, 64–67 (2013).

78. R. R. Konadhode, D. Pelluru, P. J. Shiromani, Unihemispheric Sleep: An Enigma for Current Models of Sleep-Wake Regulation, Sleep. 39 (2016)pp. 491–494.

79. G. G. Mascetti, Unihemispheric sleep and asymmetrical sleep: behavioral, neurophysiological, and functional perspectives. Nat. Sci. Sleep 8, 221–238 (2016).

80. J. J. Omoto, M. F. Keleş, B.-C. M. Nguyen, C. Bolanos, J. K. Lovick, M. A. Frye, V. Hartenstein, Visual Input to the *Drosophila* Central Complex by Developmentally and Functionally Distinct Neuronal Populations. Curr. Biol. 27, 1098–1110 (2017).

81. D. Garner, E. Kind, A. Nern, L. Houghton, A. Zhao, G. Sancer, G. M. Rubin, M. F. Wernet, S. S. Kim, Connectomic reconstruction predicts the functional organization of visual inputs to the navigation center of the *Drosophila* brain, bioRxiv (2023)p. 2023.11.29.569241.

82. B. J. Hardcastle, J. J. Omoto, P. Kandimalla, B.-C. M. Nguyen, M. F. Keleş, N. K. Boyd, V. Hartenstein, M. A. Frye, A visual pathway for skylight polarization processing in *Drosophila*. Elife 10 (2021).

83. M. Basner, Arousal threshold determination in 1862: Kohlschütter’s measurements on the firmness of sleep. Sleep Med. 11, 417–422 (2010).

84. E. Ryder, M. Ashburner, R. Bautista-Llacer, J. Drummond, J. Webster, G. Johnson, T. Morley, Y. S. Chan, F. Blows, D. Coulson, G. Reuter, H. Baisch, C. Apelt, A. Kauk, T. Rudolph, M. Kube, M. Klimm, C. Nickel, J. Szidonya, P. Maróy, M. Pal, A. Rasmuson-Lestander, K. Ekström, H. Stocker, C. Hugentobler, E. Hafen, D. Gubb, G. Pflugfelder, C. Dorner, B. Mechler, H. Schenkel, J. Marhold, F. Serras, M. Corominas, A. Punset, J. Roote, S. Russell, The DrosDel deletion collection: a *Drosophila* genomewide chromosomal deficiency resource. Genetics 177, 615–629 (2007).

85. J. D. Marshall, D. E. Aldarondo, T. W. Dunn, W. L. Wang, G. J. Berman, B. P. Ölveczky, Continuous Whole-Body 3D Kinematic Recordings across the Rodent Behavioral Repertoire. Neuron 109, 420–437.e8 (2021).

86. G. J. Berman, D. M. Choi, W. Bialek, J. W. Shaevitz, Mapping the stereotyped behaviour of freely moving fruit flies. J. R. Soc. Interface 11 (2014).

87. J. G. Todd, J. S. Kain, B. L. de Bivort, Systematic exploration of unsupervised methods for mapping behavior. Phys. Biol. 14, 015002 (2017).

88. Y. Liu, X. San Liang, R. H. Weisberg, Rectification of the Bias in the Wavelet Power Spectrum. J. Atmos. Ocean. Technol. 24, 2093–2102 (2007).

89. L. Breiman, Random Forests. Mach. Learn. 45, 5–32 (2001).

90. L. McInnes, J. Healy, J. Melville, UMAP: Uniform Manifold Approximation and Projection for Dimension Reduction, arXiv [stat.ML] (2018). http://arxiv.org/abs/1802.03426.

91. O. Friard, M. Gamba, BORIS: a free, versatile open-source event-logging software for video/audio coding and live observations. Methods Ecol. Evol. 7, 1325– 1330 (2016).

92. D. Pimentel, J. M. Donlea, C. B. Talbot, S. M. Song, A. J. F. Thurston, G. Miesenböck, Operation of a homeostatic sleep switch. Nature 536, 333–337 (2016).

93. N. Karaev, I. Rocco, B. Graham, N. Neverova, A. Vedaldi, C. Rupprecht, CoTracker: It is Better to Track Together, arXiv [cs.CV] (2023). http://arxiv.org/abs/2307.07635.

94. J. Solawetz, What is YOLOv8? The Ultimate Guide. [2024]. https://blog.roboflow.com/whats-new-in-yolov8/.

95. W. J. Joiner, A. Crocker, B. H. White, A. Sehgal, Sleep in *Drosophila* is regulated by adult mushroom bodies. Nature 441, 757–760 (2006).

